# BaiZe: A Multi-View Dynamic Framework for Simulating and Interpreting Cellular Responses Across Perturbation Contexts

**DOI:** 10.64898/2026.07.15.738608

**Authors:** Qingwu Zeng, Wenxiang Cai, Renchong Tian, Qi Wang, Dongzhan Zhou, Minting Pan, Hai Yang, Zhe Liu, Guan Ning Lin, Zhe Wang

**Affiliations:** Key Laboratory of Smart Manufacturing in Energy Chemical Process Ministry of Education, East China University of Science and Technology, Shanghai, CN; Department of Computer Science and Engineering, East China University of Science and Technology, Shanghai, CN; Shanghai Mental Health Center, Shanghai Jiao Tong University School of Medicine, School of Biomedical Engineering, Shanghai Jiao Tong University, Shanghai, China; Shanghai Gener Arfysica Intelligent Healthcare & Brain Science Research Institute, 377 Chengpu Road, Fengxian District, Shanghai 201499, China; Department of Laboratory Medicine, Ruijin Hospital,Shanghai Jiao Tong University School of Medicine, Shanghai, CN; AI for Science Center, Shanghai AI Laboratory, Shanghai, CN

**Keywords:** Single-cell perturbation modelling, conditional cellular state transition, multi-omic integration, conditional diffusion

## Abstract

Accurately predicting how cells respond to perturbations is important for understanding cellular regulation and prioritizing experimental interventions, yet existing models are often designed for specific perturbation types or biological contexts. Here we present BaiZe, a multi-view conditional state-transition framework that predicts the post-perturbation transcriptome from a control-state transcriptome together with genetic, chemical, temporal and optional chromatin-accessibility information. BaiZe models perturbation responses as context-dependent transitions between cellular states. BaiZe supports prediction across held-out cell states and genetic perturbations, unseen multi-gene combinations, chemical structures and doses, temporal stages and species contexts. Benchmarking across diverse perturbation settings demonstrates that BaiZe effectively recovers major transcriptional response programs under previously unseen conditions. Incorporating matched ATAC-seq context further improves selected state-transition predictions and enables model-based attribution of chromatin regions to response-associated genes and pathways. BaiZe also supports few-shot transfer of perturbation responses from human to mouse cellular systems and connects predicted transcriptomes to candidate morphology projections. To facilitate interpretation and use, BaiZe-Agent organizes response genes, pathways, chromatin evidence, cross-species predictions and projected phenotypes into traceable, queryable perturbation records. Together, BaiZe provides a broadly applicable framework for predicting and interpreting context-dependent cellular responses and for prioritizing hypotheses across diverse perturbation settings.

## Introduction

Single-cell perturbation assays connect controlled interventions to molecular responses, providing a powerful basis for functional genomics, disease-mechanism studies, therapeutic target discovery and phenotypic screening. Pooled CRISPR screens with single-cell transcriptomic readouts resolve context-dependent genetic effects at scales that are inaccessible to conventional one-gene-at-a-time experiments [1–4], while multiplexed chemical profiling extends this principle across compounds, doses and cellular backgrounds [5]. Harmonized data resources and end-to-end analysis ecosystems have further improved the comparison and reuse of perturbation experiments [6, 7]. Yet experimental coverage remains sparse because the space defined by genes and gene combinations, compounds and doses, cellular backgrounds, temporal states and species expands far more rapidly than feasible screening capacity. Computational models are therefore needed to predict unmeasured responses, prioritize informative interventions and identify conditions in which additional experiments would be most valuable. These goals are central to the virtual-cell agenda, which seeks predictive and queryable connections among molecular state, intervention and phenotype [8, 9].

Existing computational approaches have advanced perturbation-response modelling along several complementary directions. Latent-space state-translation models, including scGen and trVAE, learn mappings between control and perturbed cell distributions and support response transfer across cellular contexts [10, 11]. Compositional perturbation autoencoders extend this formulation to structured conditions: CPA represents perturbation combinations, dose and context, and chemCPA incorporates molecular structure to support extrapolation to unseen compounds [12, 13]. CellOT models treatment-induced distributional changes through neural optimal transport, while biolord separates known attributes from latent cellular variation [14, 15]. GEARS incorporates gene relationships to predict unmeasured genetic combinations [16], and chemical-response models increasingly use compound representations, transfer learning and biological priors to address novel compounds, unseen cellular backgrounds and data-limited settings [17–19]. Large-scale single-cell pretraining in Geneformer, scGPT and scFoundation supports downstream network and perturbation tasks [20–22], while universal cell embedding models emphasize transferable representations across datasets and species [23]. Collectively, these studies show that intervention identity, initial cellular state and biological structure can improve prediction under unmeasured conditions.

Despite this progress, perturbation-response modelling remains fragmented across intervention types and biological settings. Genetic perturbations, multi-gene combinations, chemical structure and dose, temporal transitions, chromatin-conditioned responses and cross-species transfer are generally addressed through separate models or evaluation protocols. Differences in assay design, control quality, perturbation efficiency and response strength further limit direct comparison across datasets [6]. Recent benchmarks have also shown that model rankings depend strongly on data partitioning, metric selection and baseline construction [24–27]. Whole-transcriptome similarity can be dominated by shared cellular identity and systematic variation, producing favourable scores even when key response genes, effect magnitudes or expression directions are recovered poorly [28]. Generalization to an unseen cell state, compound, gene combination or species also represents a distinct predictive problem. Zero-shot transfer evaluates whether information learned in a source domain can support prediction in a new target domain, while few-shot adaptation measures how effectively limited target-domain observations calibrate systematic differences. Robust evaluation therefore requires response-focused metrics alongside global similarity, explicit assessment of expression direction and magnitude, and clear separation between predictive associations and experimentally established causal effects [29]. Uncertainty-aware approaches such as GPerturb provide a complementary means of characterizing confidence in estimated perturbation effects [30].

Accurate response prediction also depends on biological context extending beyond the baseline transcriptome. RNA profiles describe the current transcriptional state, while ATAC-seq measures accessible chromatin and provides complementary information about regulatory potential [31]. ArchR and Signac support scalable analysis of chromatin accessibility and peak-to-gene relationships, while MultiVI and totalVI illustrate probabilistic integration across single-cell modalities [32–35]. Paired profiling studies have shown that accessibility changes can precede transcriptional changes during differentiation [36], and joint CRISPR multi-omic screens increasingly connect genetic perturbations with enhancer activity, chromatin state and downstream transcriptional responses [37, 38]. Phenotypic information provides another complementary view: transcriptome-guided image models can project predicted molecular states into morphology space [39], while experimental profiling methods such as Cell Painting establish image-based representations of cellular phenotype [40]. Cross-species prediction introduces an additional layer of biological structure because conserved responses coexist with differences in orthology, paralogy, cellular composition and species-specific regulation. Protein-informed and graph-based representations can improve biological alignment across species, but limited target-domain observations may still be needed to calibrate species-specific effects [41, 42]. Together, these considerations motivate a framework that can incorporate multiple condition types, distinguish zero-shot transfer from few-shot adaptation and retain explicit links between each prediction and its supporting molecular evidence.

Here we present BaiZe, a multi-view conditional cellular state-transition framework for predicting context-dependent responses to genetic, chemical, temporal, regulatory and cross-species conditions. BaiZe represents each prediction as a transition from a control-state transcriptome to a post-perturbation transcriptome. A cellular-background encoder captures the initial molecular state, condition encoders represent the applied intervention and available contextual information, and a conditional diffusion model generates the corresponding expression change as a transcriptomic residual. This common formulation supports held-out cellular-state prediction, genetic perturbations, unseen multi-gene combinations, drug structure and dose modelling, temporal projection and human-to-mouse perturbation transfer under zero-shot and few-shot settings. The cross-species immune analysis was motivated by functional CRISPR studies showing that chromatin regulators influence T-cell state and persistence in mouse tumour-infiltrating lymphocytes and human CD8+ T cells [52, 54]. Matched ATAC context provides an optional regulatory view, with peak-level attribution linking model-used chromatin regions to candidate genes and pathways. Predicted transcriptomes can also be projected into morphology space to obtain morphology representations associated with the predicted molecular states. BaiZe-Agent complements the predictive framework by organizing predicted response genes, pathways, chromatin evidence, cross-species patterns and morphology projections into traceable records that can be explored through natural-language queries.

We evaluate BaiZe using whole-transcriptome and response-focused criteria, including Top20 Delta Pearson, mean-squared error and opposite-direction rate, together with task-matched baselines and biologically structured held-out settings. Across the evaluated tasks, BaiZe recovered major perturbation-associated transcriptional programs in held-out cellular and intervention settings and supported human-to-mouse transfer under zero-shot and few-shot evaluation settings. Matched chromatin information improved selected temporal-state predictions and provided candidate regulatory evidence through peak-level attribution, while limited target-species observations improved the calibration of cross-species predictions. By combining conditional state-transition modelling, multi-view cellular context and traceable result exploration, BaiZe provides a common computational framework for studying cellular responses across diverse perturbation settings and prioritizing experimentally testable hypotheses.

## Results

### BaiZe Establishes a Multi-View Conditional State-Transition Framework

Cellular responses to perturbations can be represented as state transitions conditioned on the initial cellular context and the applied intervention [10, 12, 14, 16]. We formalized this process as conditional cellular state transition modelling and developed BaiZe to predict perturbation-induced transcriptomic responses. BaiZe receives a control-state transcriptome, combines it with a specified perturbation context and predicts the expression change that defines the post-perturbation transcriptome. The resulting prediction can then be analysed through response-gene ranking, pathway enrichment, regulatory attribution, cross-species transfer and morphology projection.

The core architecture of BaiZe consists of a cellular-background encoder, a condition encoder and a conditional diffusion decoder. The cellular-background encoder maps the control-state RNA profile into a latent representation of the starting cell state. This representation captures cell identity, baseline transcriptional programs and pre-existing cell-to-cell variation. The condition encoder transforms the intervention into a shared latent condition, including genetic perturbations, combinatorial targets, drug structure, dose, time or state labels, cell-context information and optional ATAC-derived regulatory context. The diffusion decoder then generates a perturbation-specific transcriptomic residual through iterative denoising. Adding this residual to the control expression profile yields the predicted post-perturbation transcriptome (Fig. 1a). This organization keeps the model architecture unified while allowing each task to provide its own biological condition and evaluation setting.

**Figure 1.**
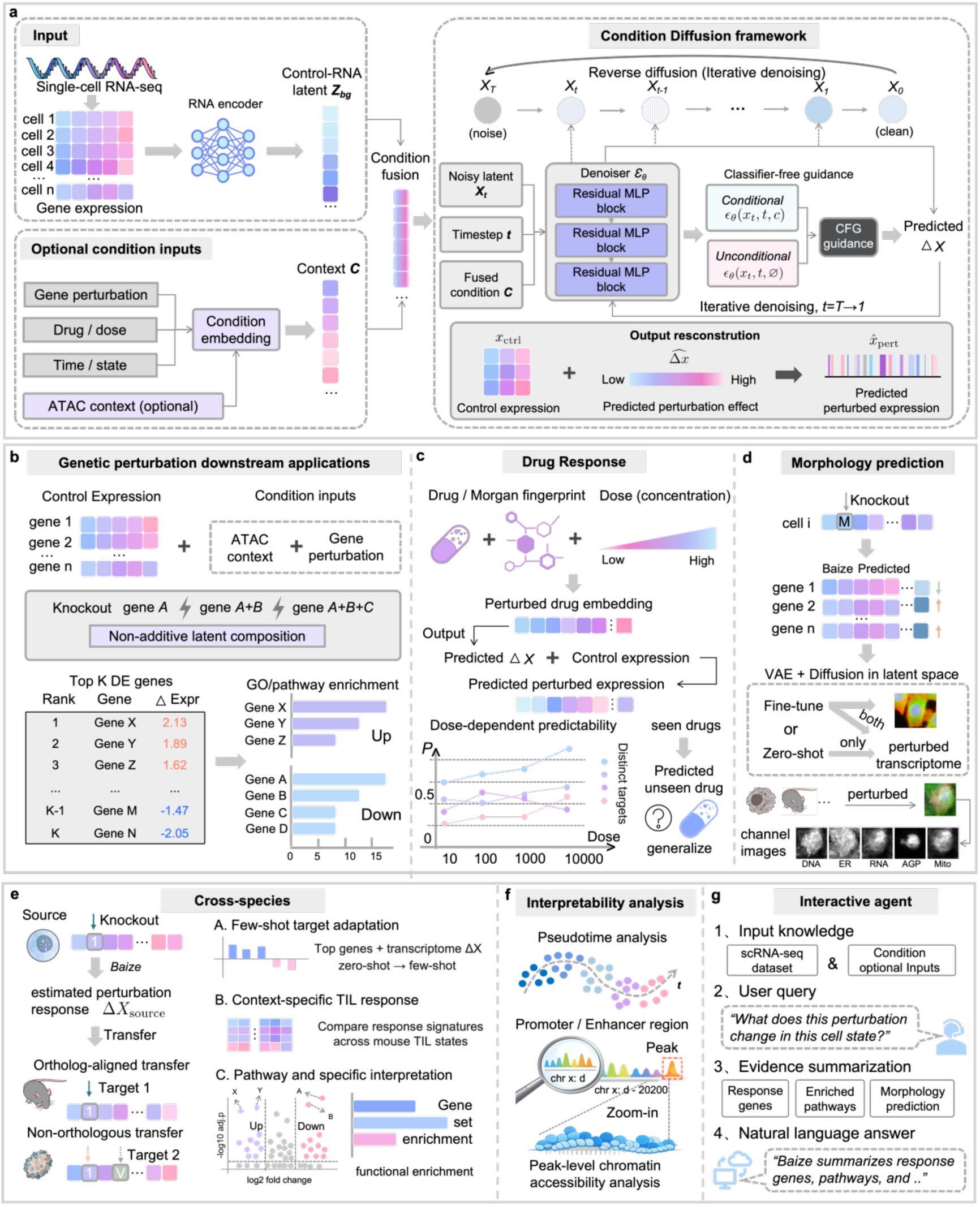
Model architecture and downstream applications of BaiZe. (a) encoding control-state single-cell RNA expression as the cellular background and encoding gene perturbation, drug/dose, time/state and optional ATAC context as conditions; (b) predicting single-gene, multi-gene and combinatorial perturbation responses, followed by response gene ranking and pathway enrichment analysis; (c) using drug molecular fingerprints and dose information to predict dose-dependent transcriptional responses and evaluate generalization to unseen drugs; (d) using the predicted post-perturbation transcriptome as input for transcriptome-guided morphology projection; (e) transferring perturbation-response knowledge across species through ortholog-aligned gene spaces and supporting zero-shot, few-shot and target-species-specific response analysis; (f) linking ATAC-informed perturbation predictions to pseudotime changes, promoter/enhancer regions and peak-level chromatin accessibility; (g) summarizing response genes, enriched pathways, morphology projection and other evidence through natural-language user queries.

First, BaiZe learns perturbation responses in the residual-expression space. The model estimates the expression change associated with a given intervention and adds this shift to the control-state profile to reconstruct the predicted post-perturbation state. This formulation anchors the prediction to the starting cellular context and focuses the generative process on intervention-associated transcriptional changes. The resulting expression changes provide a basis for response-gene ranking, comparison of single-and multi-gene perturbation responses, and downstream functional interpretation (Fig. 1b).

Second, BaiZe represents heterogeneous perturbation contexts through a unified condition interface. Genetic targets, combinatorial perturbations, drug structure, dose and temporal labels are encoded as condition signals and integrated with the same cellular-background representation. In the drug-response module, BaiZe uses Morgan fingerprints and dose information to predict dose-dependent transcriptional changes and to evaluate generalization to unseen drugs or unseen cellular backgrounds (Fig. 1c). In the temporal module, target time or state labels provide the condition for projecting cells towards later transcriptional states. This shared generative formulation supports perturbation, treatment and temporal prediction tasks while preserving task-specific inputs.

Third, BaiZe connects transcriptional response prediction with additional cellular views. Predicted post-perturbation transcriptomes can serve as molecular inputs for morphology projection, generating multi-channel phenotype hypotheses across DNA, ER, RNA, AGP and mitochondrial signals (Fig. 1d). When paired RNA+ATAC data are available, chromatin-accessibility context can be incorporated into the condition representation to refine response prediction. The ATAC-informed outputs can then be analysed to identify promoter, enhancer and peak-level accessibility signals associated with improved prediction of selected response genes, providing candidate regulatory evidence for further inspection (Fig. 1f). For cross-species analysis, ortholog-aligned gene spaces support the transfer of perturbation information from human cells to mouse tumour-infiltrating T-cell systems, enabling state-specific response prediction in the target species (Fig. 1e).

We further implemented BaiZe-Agent to help users explore simulated perturbation results through natural-language queries (Fig. 1g). After a perturbation is simulated, the model output often spans several layers of evidence, including predicted expression changes, response-gene rankings, regulatory attribution and projected phenotypes. BaiZe-Agent organizes these results into an interactive query workflow. Each BaiZe simulation therefore yields more than a predicted transcriptome. It produces an interpretable perturbation record that links the estimated expression change to candidate response genes, regulatory evidence and downstream phenotype hypotheses.

### Model generalizes across perturbation settings and supports morphology projection

We next asked whether BaiZe could predict perturbation responses beyond the conditions used for training. We first tested whether BaiZe could predict perturbation responses in an unseen cellular state. Human embryo 10x Multiome data provide a suitable setting for this task because they contain multiple developmental cell states with paired RNA and ATAC profiles [43]. We held out the HYPO/VE state during training and used it only for testing, while AME and EXMC were used to learn perturbation-response patterns (Fig. 2a).

**Figure 2.**
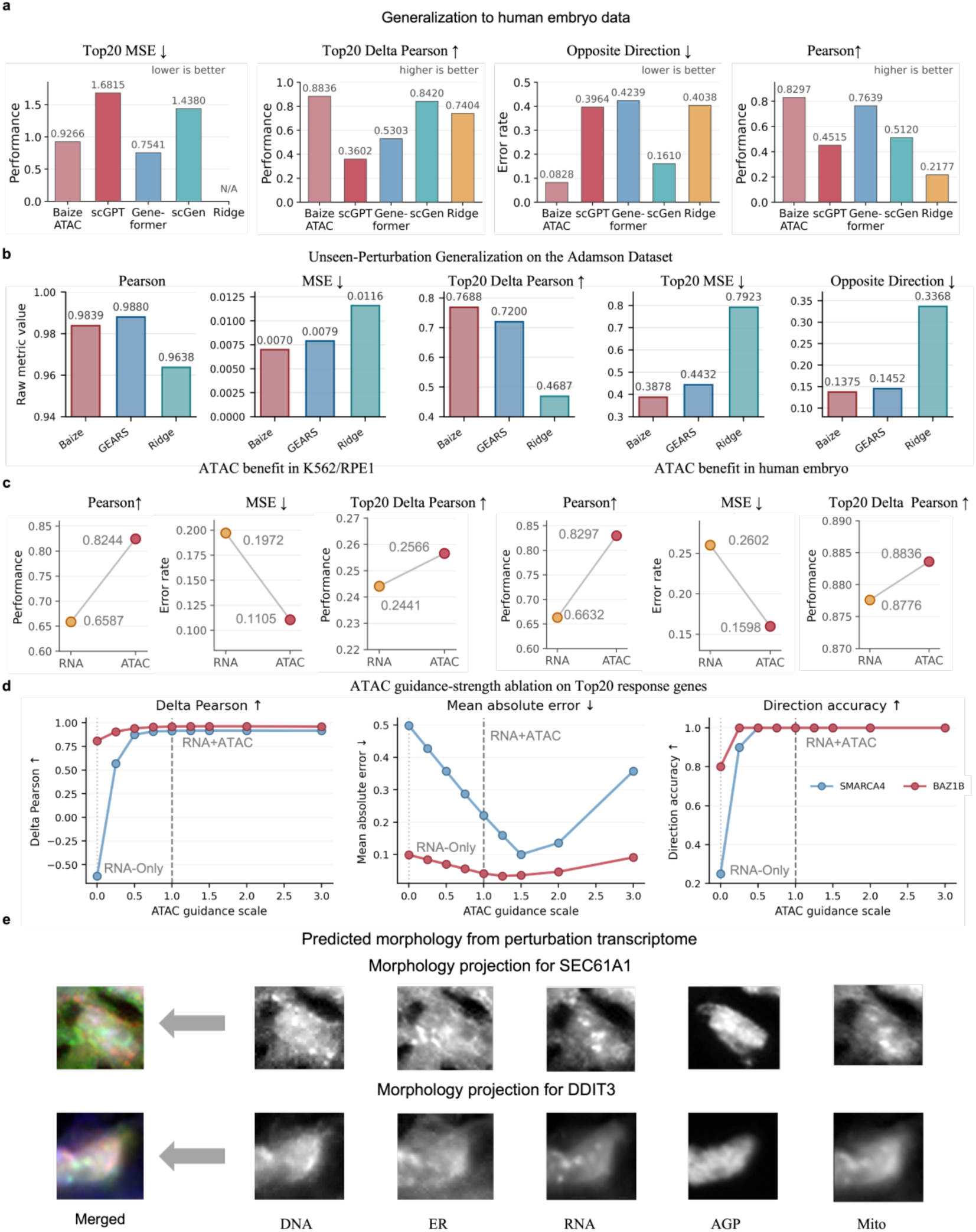
Generalization to unseen perturbation settings and morphology projection. (a) evaluating BaiZe on a held-out HYPO/VE cell-type task in human embryo 10x Multiome data and comparing it with scGPT, Geneformer, scGen and Ridge; (b) testing unseen-perturbation generalization on the Adamson dataset and comparing BaiZe with GEARS and Ridge across whole-gene and Top20 response-gene metrics; (c) comparing RNA-only and RNA+ATAC settings in K562/RPE1 and human embryo data to assess the contribution of ATAC context; (d) varying the ATAC guidance scale to evaluate its effect on Top20 Delta Pearson, mean absolute error and direction accuracy; (e) projecting BaiZe-predicted post-perturbation transcriptomes for *SEC61A1* and *DDIT3* into five-channel morphology space using the pretrained MorphDiff G2I model.

We compared BaiZe with scGPT [21], Geneformer [20], scGen [10] and Ridge [25], covering foundation-model, generative and linear baselines. BaiZe achieved the highest whole-transcriptome Pearson correlation (0.8297), the highest Top20 Delta Pearson correlation (0.8836) and the lowest opposite-direction rate (0.0828) in the held-out HYPO/VE state (Fig. 2a). The Top20 Delta Pearson correlations of scGPT [21], Geneformer [20], scGen [10] and Ridge were 0.3602, 0.5303, 0.8420 and 0.7404, and their opposite-direction rates were 0.3964, 0.4239, 0.1610 and 0.4038, respectively. BaiZe also achieved a Top20 MSE of 0.9266, lower than scGPT and scGen, while Geneformer obtained a lower value on this metric. These results indicate that BaiZe more consistently recovered the dominant response-gene trends and directions of expression change in a cellular state excluded from training.

We next tested unseen perturbation generalization on the Adamson Perturb-seq dataset, which contains CRISPR perturbations related to ER stress and the unfolded protein response (Fig. 2b) [2]. BaiZe reached a whole-transcriptome Pearson correlation correlation of 0.9839, close to GEARS [16] (0.9880), and showed stronger performance on response-focused metrics. Its Top20 Delta Pearson correlation was 0.7688, compared with 0.7200 for GEARS [16] and 0.4687 for Ridge. Its Top20 MSE was 0.3878, lower than GEARS (0.4432) and Ridge (0.7923), and its opposite-direction rate was 0.1375. BaiZe also achieved the lowest whole-gene MSE (0.0070). These results suggest that BaiZe can recover major transcriptional responses for held-out perturbation conditions.

We then assessed the contribution of chromatin-accessibility context using RNA-only and RNA+ATAC versions of BaiZe in K562/RPE1 and human embryo tasks (Fig. 2c) [37,43]. The two versions used the same model structure and training strategy, with ATAC latent context added in the RNA+ATAC setting. In K562/RPE1, ATAC increased Pearson correlation from 0.6587 to 0.8244 and reduced MSE from 0.1972 to 0.1105. In the human embryo task, Pearson correlation increased from 0.6632 to 0.8297 and MSE decreased from 0.2602 to 0.1598. Top20 Delta Pearson showed smaller gains, increasing from 0.2441 to 0.2586 in K562/RPE1 and from 0.8776 to 0.8836 in human embryo. An ATAC guidance-scale ablation further showed that calibrated ATAC guidance improved response recovery for *SMARCA4* and *BAZ1B*, while overly strong guidance reduced or plateaued performance in several metrics (Fig. 2d). Component-level ablations supported the same conclusion, with ATAC context improving delta-based response recovery and the addition of gene prior and diffusion residual further increasing perturbation-level Top20 Delta Pearson in the ATAC-informed setting (Supplementary Table S3).

Finally, we examined whether BaiZe-predicted post-perturbation transcriptomes could be projected into morphology space (Fig. 2e). Predicted transcriptomes for *SEC61A1* and *DDIT3* perturbations were provided to the pretrained MorphDiff [39] gene-to-image (G2I) model, which generated projections across DNA, endoplasmic reticulum (ER), RNA, actin/Golgi/plasma membrane (AGP) and mitochondrial (Mito) channels. These images illustrate candidate morphological readouts associated with the predicted molecular responses. Because paired microscopy images were unavailable, the outputs were interpreted as transcriptome-derived candidate morphology projections, with no image-level reconstruction validation.

### BaiZe predicts held-out multi-gene perturbation responses and captures combination-specific patterns

To assess whether BaiZe could generalize from observed perturbations to unseen multi-gene combinations, we constructed held-out prediction tasks using the Norman single-cell CRISPR perturbation dataset [4]. BaiZe represents multiple target genes together as a single perturbation condition, allowing the model to handle combinations containing different numbers of genes. We compared this approach with a simple baseline that predicts a combination by directly adding the expression changes caused by the corresponding single-gene perturbations. Test combinations were grouped into three settings according to how much information about their constituent genes was available during training. The most difficult setting contained combinations whose genes had the least training coverage, whereas the easiest setting contained genes that had been observed individually or in other combinations, but not in the specific test combination. BaiZe achieved higher Top20 Delta Pearson correlation and lower Top20 mean squared error across these settings, with the largest improvement in the most challenging group (Fig. 3a and Supplementary Table S4). Gene-level analyses also showed that BaiZe more accurately recovered major response-gene patterns and their directions of change (Supplementary Fig. S1). These results indicate that BaiZe can use previously observed perturbations to predict transcriptional responses to unseen gene combinations.

**Figure 3.**
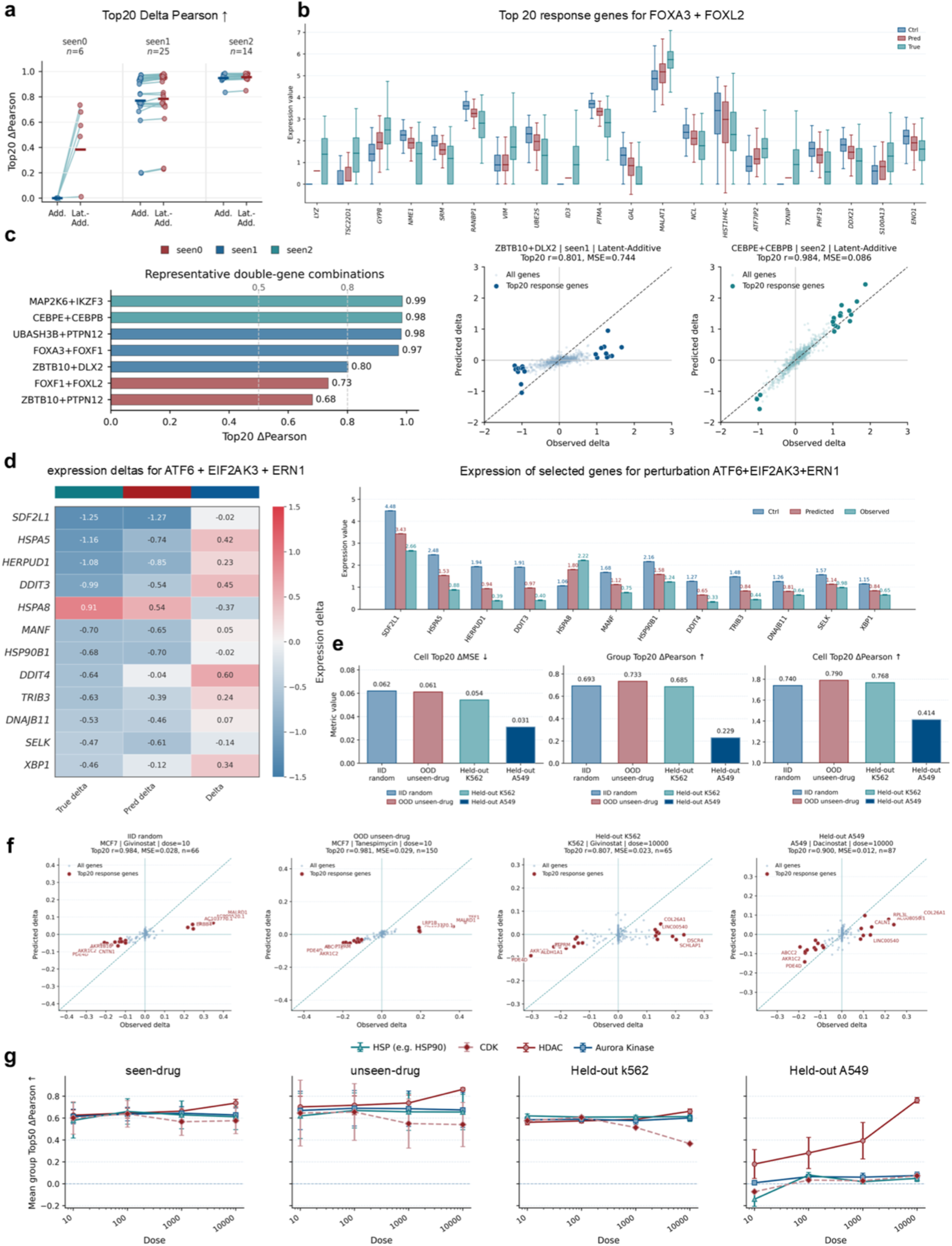
Prediction of combinatorial genetic perturbations and drug-induced transcriptional responses by BaiZe. (a) comparing Expression-space addition and Latent-space conditioning strategies across held-out combinatorial perturbation settings with different levels of gene overlap; (b) evaluating predicted expression distributions of Top20 response genes for the *FOXA3+FOXL2* double-gene perturbation; (c) benchmarking representative held-out double-gene combinations and comparing observed and predicted expression changes for Top20 response genes; (d) testing BaiZe on the *ATF6+EIF2AK3+ERN1* triple-gene perturbation and comparing observed, predicted and residual-target expression changes; (e) evaluating drug-response prediction across seen-drug, unseen-drug, held-out K562 and held-out A549 settings; (f) showing representative drug-response predictions by comparing observed and predicted expression changes across different generalization settings; (g) stratifying group-level Top50 Delta Pearson by drug mechanism and dose to assess dose-dependent predictability.

For representative double-gene perturbations, BaiZe recovered the major response-gene trends with high accuracy. For *FOXA3+FOXL2*, the predicted expression distributions of the Top20 response genes were closer to the observed post-perturbation state and were distinguishable from those of control cells (Fig. 3b). Across several held-out double-gene combinations, predictions based on a joint representation of the perturbation targets maintained high correlations for the major response genes. Top20 Delta Pearson correlations were close to or above 0.97 for MAP2K6+IKZF3, *CEBPE+CEBPB*, *UBASH3B+PTPN12* and *FOXA3+FOXF1*, whereas the corresponding values for *ZBTB10+DLX2* and *ZBTB10+PTPN12* were 0.801 and 0.68, respectively, indicating variation in predictability across gene combinations (Fig. 3c). Scatter plots further showed close agreement between predicted and observed changes for the Top20 response genes in *ZBTB10+DLX2* and *CEBPE+CEBPB*, with Top20 Delta Pearson correlations of 0.801 and 0.984. Compared with directly adding the effects of individual gene perturbations, joint modelling achieved higher predictive correlations for multiple combinations, supporting its ability to recover combination-specific transcriptional responses.

To extend the evaluation to a higher-order gene combination, we held out the *ATF6+EIF2AK3+ERN1* triple-gene perturbation from the Adamson Perturb-seq dataset [2] for testing (Fig. 3d). This combination is associated with endoplasmic reticulum (ER) stress and the unfolded protein response (UPR). BaiZe recovered the observed direction of change for several major response genes, including *SDF2L1, HSPA5, HERPUD1, DDIT3, HSP90B1, MANF* and *DNAJB11*. The heatmap showed that the predictions captured the main response patterns, while the selected-gene expression levels shifted from the control state towards the observed post-perturbation state. Gene-level expression means and changes for representative UPR response genes further supported this directional agreement (Supplementary Table S5). These results indicate that BaiZe can apply response patterns learned from observed single- and double-gene perturbations to predict the transcriptional response of an unseen triple-gene combination.

### Chemical structure and dose modelling predicts drug perturbation responses

Drug-response prediction can help prioritize candidate compounds and doses before large-scale experimental screening, but the task is complicated by the dependence of transcriptional responses on dose and cellular context. To assess whether BaiZe could use chemical structure to predict unmeasured drug responses, we evaluated the model on the Sci-Plex single-cell drug perturbation dataset [5]. BaiZe received control-state RNA, Morgan fingerprints derived from SMILES strings [53] and treatment dose as inputs, and predicted transcriptional changes at 10, 100, 1,000 and 10,000 nM. Drug names, molecular targets and mechanism-of-action annotations were used only for sample annotation and downstream stratification.

We evaluated four settings: a random cell split within observed experimental conditions, an unseen-drug split in which test compounds were excluded from training, and held-out K562 and A549 cell-context splits (Fig. 3e). Performance was assessed at both the single-cell level and the group level after aggregation by drug, dose and cellular context. In the random, unseen-drug and held-out K562 settings, group-level Top20 Delta Pearson correlations were 0.693, 0.733 and 0.685, respectively, while the corresponding single-cell values were 0.740, 0.790 and 0.768. Single-cell Top20 delta MSE values were 0.062, 0.061 and 0.054, indicating that BaiZe retained the ability to recover major response-gene trends even when the test compounds were absent from training.

Generalization across cellular contexts was more variable. In the held-out A549 setting, group-level and single-cell Top20 Delta Pearson correlations decreased to 0.229 and 0.414, respectively, indicating that drug-response patterns learned from other cellular backgrounds did not fully recover the major responses in A549 cells. The relatively low Top20 delta MSE in this setting was not accompanied by a comparable increase in correlation, suggesting that the model tended to predict changes of smaller magnitude. Complete results across random, unseen-drug and held-out cell-context settings, including the weaker held-out MCF7 result, are reported in Supplementary Table S6.

Representative examples further illustrated variation across prediction settings (Fig. 3f). In MCF7 cells, the Top20 Pearson correlation reached 0.984 for low-dose Givinostat in the random split and 0.981 for the unseen drug Tanespimycin. In the held-out K562 and A549 settings, high-dose Givinostat and high-dose Dacinostat achieved Top20 Pearson correlations of 0.807 and 0.900, respectively. Stratification by drug mechanism and dose revealed distinct group-level Top50 Delta Pearson trends across cellular backgrounds (Fig. 3g). Histone deacetylase (HDAC) inhibitors showed improved performance with increasing dose in the held-out A549 setting, whereas Aurora kinase, heat-shock protein (HSP/HSP90) and cyclin-dependent kinase (CDK) compounds remained less predictable. Additional analyses across molecular targets, pathways and recurrent response genes similarly showed that drug-response predictability varied with mechanism and cellular context (Supplementary Fig. S2).

### Regulatory attribution identifies mechanisms underlying perturbation-response correction

The value of a multi-omic perturbation-response model depends not only on predictive accuracy, but also on whether the contribution of additional molecular modalities can be interpreted. RNA expression reflects the current transcriptional state of a cell, whereas ATAC-seq captures chromatin accessibility and may reveal regulatory potential that is not fully represented in the transcriptome [31]. To examine whether this regulatory context could explain improvements in perturbation-response prediction, we compared RNA-only and RNA+ATAC models and defined ATAC-corrected genes as response genes for which the inclusion of ATAC reduced prediction error while preserving the observed direction of change. Ablation analysis showed that RNA+ATAC improved perturbation-level Delta Pearson and Top20 Delta Pearson, whereas masking promoter-associated peaks reduced these gains (Supplementary Table S7). This result suggests that promoter accessibility provides useful information for determining whether promoter accessibility provides information associated with transcriptional responsiveness.

Under *SMARCA4* and *BAZ1B* perturbations, ATAC information corrected a subset of response genes that appeared sensitive to chromatin-regulatory context (Fig. 4a). *SMARCA4* encodes the catalytic ATPase subunit of the SWI/SNF chromatin-remodelling complex, and its perturbation can alter nucleosome organization and local DNA accessibility, thereby affecting downstream transcriptional programs [44]. In this setting, RNA+ATAC reduced prediction errors for genes including *NRG1*, *ERBB4*, *ANK2* and *LAMC1*, which are involved in intercellular signaling, membrane organization and extracellular-matrix interactions [47]. *BAZ1B* is an ATP-dependent chromatin regulator involved in transcriptional control and neurodevelopmental differentiation [45, 46]. In our analysis, the genes corrected after incorporating ATAC context included *MALAT1, CACNA1A, AFF3* and *NALADL2*, with functional annotations related to RNA regulation, ion-channel activity and cellular-state maintenance. Top20 Delta Pearson increased from −0.723 to 0.915 for *SMARCA4* and from −0.783 to 0.890 for *BAZ1B*, accompanied by lower MSE in both cases (Supplementary Table S8).

**Figure 4.**
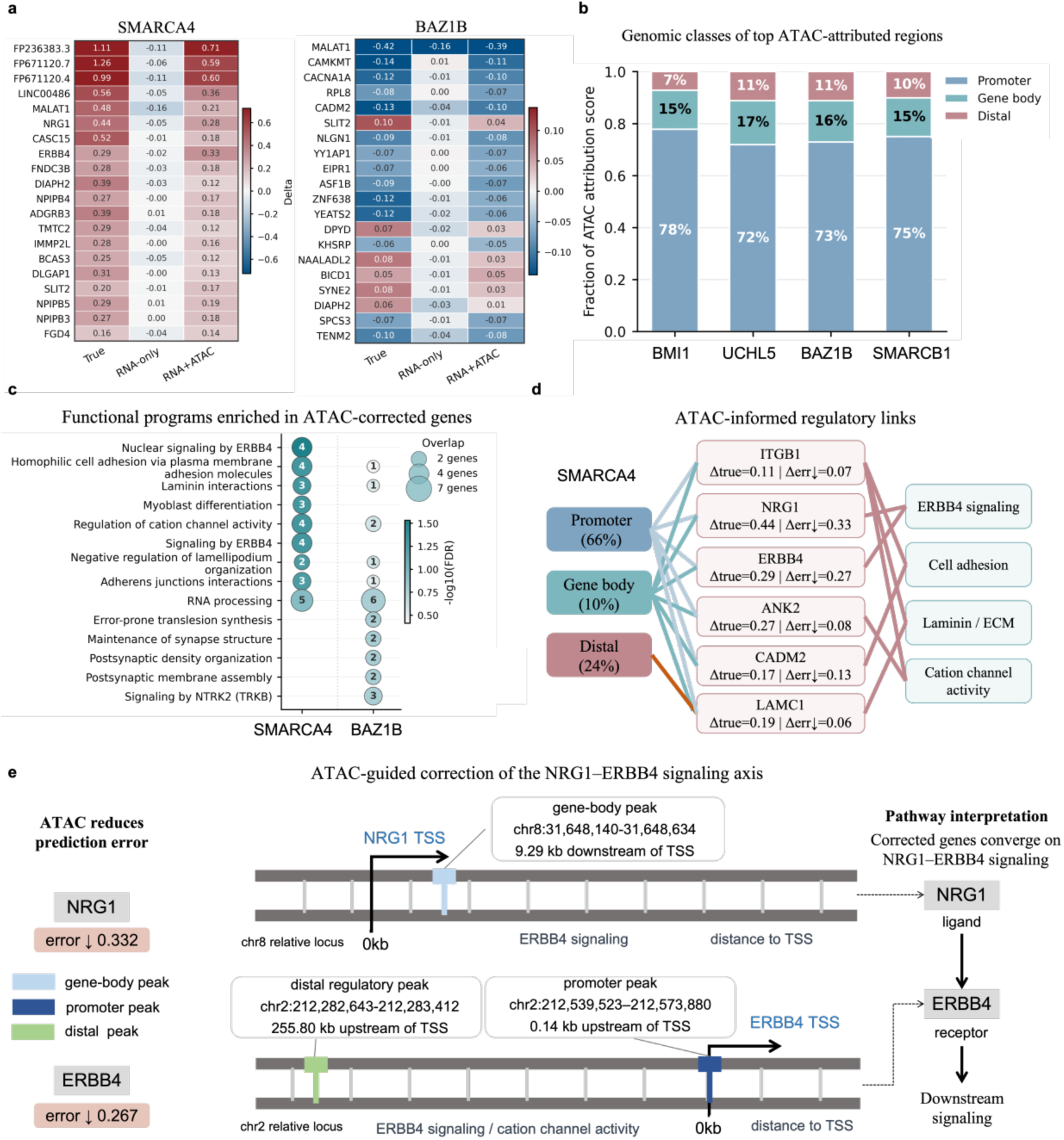
Regulatory attribution of ATAC-guided perturbation-response correction. (a) comparing observed, RNA-only predicted and RNA+ATAC predicted expression changes for representative ATAC-corrected genes under *SMARCA4* and *BAZ1B* perturbations; (b) annotating top ATAC-attributed regions into promoter, gene body and distal genomic classes; (c) identifying functional programs enriched in ATAC-corrected genes, including ERBB4 signaling, cell adhesion, laminin/ECM interaction and cation channel activity; (d) linking high-contribution genomic regions, corrected genes and functional modules in the *SMARCA4* perturbation; (e) showing an *NRG1–ERBB4* signaling-axis example in which ATAC-attributed peaks near *NRG1* and *ERBB4* are associated with reduced prediction error.

To identify the genomic sources associated with these corrections, we mapped highly attributed ATAC signals back to specific chromatin regions. Across perturbations including *BMI1, UCHL5, BAZ1B* and *SMARCB1*, the highest-attribution regions were concentrated near promoters, with additional contributions from gene-body and distal regions. Across eight perturbations, promoter-proximal regions accounted for approximately 72%–78% of the top attributed regions, while the remaining signals were distributed across gene-body and distal regions (Fig. 4b and Supplementary Table S9). Because promoter accessibility is closely related to transcription initiation, its dominant contribution suggests that the model relied strongly on regulatory information reflecting gene activation potential. Gene-body accessibility may capture ongoing transcription and local chromatin organization, whereas distal regions may contain enhancers or other long-range regulatory elements.

Functional enrichment analysis further showed that ATAC-corrected genes under *SMARCA4* perturbation were associated with *ERBB4* signaling, cell adhesion, laminin interactions, adherens junctions and cation-channel activity (Fig. 4c). Together, these programs point to changes in intercellular communication, membrane signaling, extracellular-matrix attachment and structural remodelling, consistent with broader cell-state changes following chromatin-remodelling perturbation. Enrichment signals under *BAZ1B* perturbation were weaker and were mainly related to RNA processing and neuronal structural organization. These associations were therefore interpreted as supporting trends that require further validation.

The NRG1–ERBB4 signaling axis under *SMARCA4* perturbation provides a representative example of how chromatin regions, response genes and functional programs can be connected (Fig. 4d,e). *NRG1* encodes a ligand of the *ERBB4* receptor family, whereas *ERBB4* encodes a receptor tyrosine kinase involved in intercellular signaling, differentiation and tissue-state regulation [47]. Incorporating ATAC reduced the prediction errors of *NRG1* and *ERBB4* by 0.332 and 0.267, respectively. Highly attributed signals included a gene-body peak near *NRG1*, a promoter-proximal peak near the *ERBB4* transcription start site and a distal peak located approximately 255.8 kb from the *ERBB4* transcription start site. Corrected genes including *NRG1, ERBB4, ITGB1, ANK2, CADM2* and *LAMC1* were further linked to *ERBB4* signaling, cell adhesion, laminin and extracellular-matrix interactions, and cation-channel activity. Fine-grained peak-to-gene analysis identified corresponding attribution patterns across promoter, gene-body and distal regions (Supplementary Fig. S3). These results define a testable regulatory hypothesis, while the inferred peak-to-gene links remain model-based associations that require experimental validation.

### BaiZe enables cross-species prediction of perturbation responses across mouse TIL states

Cross-species perturbation prediction provides a route for transferring experimentally derived knowledge from one species to another, with potential applications in disease modelling and experimental hypothesis prioritization. The main challenge is that human and mouse cells differ in control-state RNA profile programs, cellular composition and regulatory context, causing the same perturbation to produce distinct responses in the target species. To evaluate the cross-species transferability of BaiZe, we constructed a prediction task using perturbation-response data from human CD8+ T cells and mouse tumour-infiltrating lymphocytes derived from previously published CRISPR studies [52, 54]. Human and mouse datasets were aligned through one-to-one orthologs, yielding a shared space of 15,520 genes. The target dataset contained six mouse TIL states: C1-S1pr1, C2-ISG, C3-41BB, C4-ProgEx, C5-TermEx and C6-Cycling. In the zero-shot setting, no post-perturbation mouse RNA profiles were used for training. BaiZe used human *ARID1A* and *PDCD1* perturbation data together with unperturbed mouse RNA and target-species ATAC context to predict mouse perturbation responses. We compared BaiZe with an average-response transfer baseline, scGPT [21], UCE [23] and an RNA-only version of BaiZe, covering direct transfer, pretrained single-cell models and a within-framework ablation. BaiZe (RNA+ATAC) achieved the highest mean Top20 Delta Pearson correlation of 0.4348 and the lowest Top20 MSE (Fig. 5a), indicating that the baseline transcriptional state and chromatin context of the target species provided useful calibration for cross-species response prediction (Supplementary Table S10).

**Figure 5.**
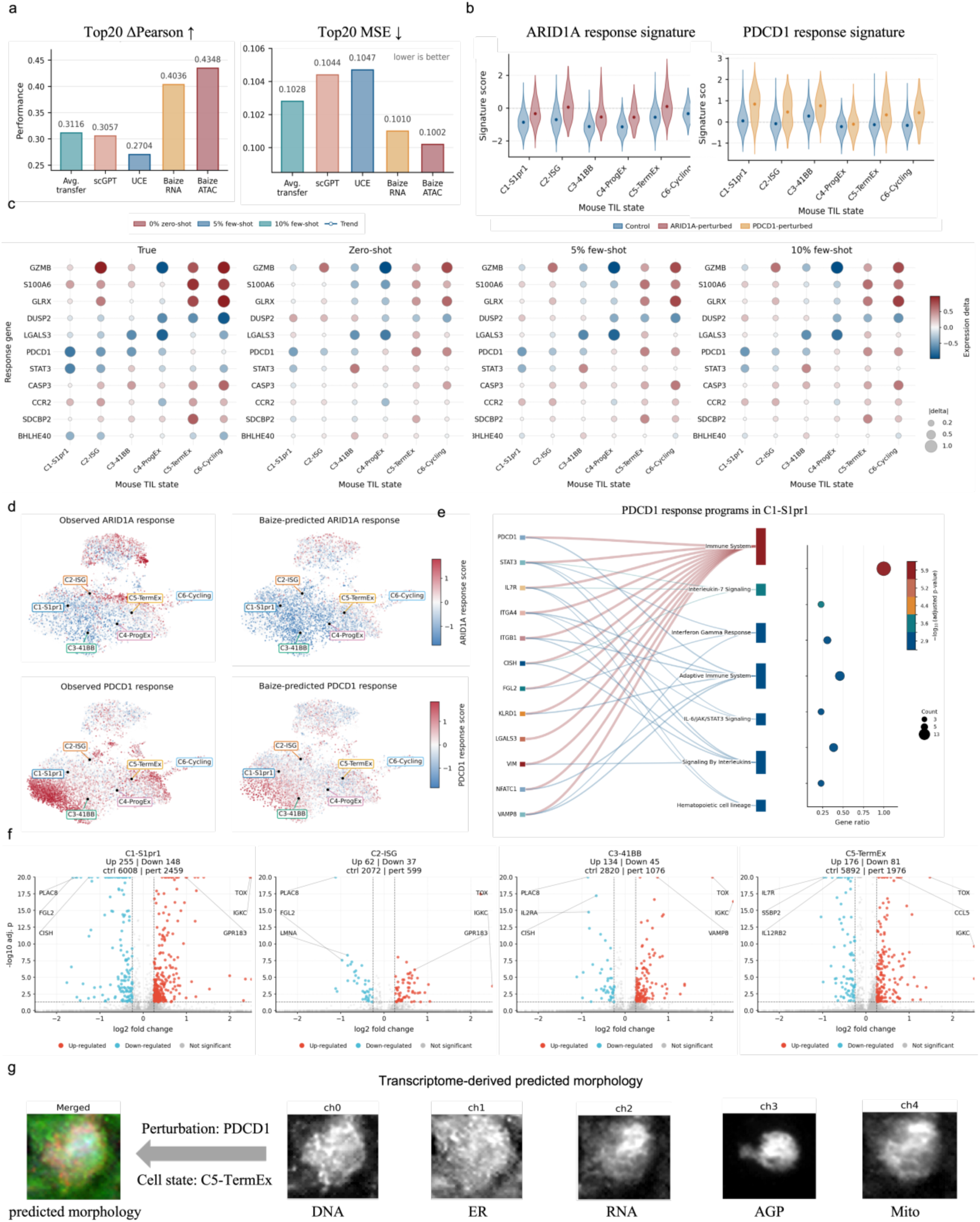
Cross-species perturbation-response prediction across mouse TIL states. (a) comparing zero-shot cross-species prediction performance across methods using Top20 Delta Pearson and Top20 MSE; (b) scoring *ARID1A* and *PDCD1* perturbation-response signatures across mouse TIL states; (c) comparing observed, zero-shot, 5% few-shot and 10% few-shot *ARID1A* responses for representative genes across TIL states; (d) projecting observed and BaiZe-predicted *ARID1A* and *PDCD1* response signatures onto the mouse TIL UMAP space; (e) linking *PDCD1* response genes in C1-S1pr1 cells to immune-related pathways; (f) identifying context-specific *PDCD1* response genes across different mouse TIL states using differential expression analysis; (g) projecting the BaiZe-predicted *PDCD1* post-perturbation transcriptome in the C5-TermEx state into five-channel morphology space using the pretrained MorphDiff G2I model.

The predicted responses to *ARID1A* and *PDCD1* perturbations were distributed differently across mouse TIL states (Fig. 5b). *ARID1A* response signals were stronger in C2-ISG, C5-TermEx and C6-Cycling cells, whereas *PDCD1* responses showed greater state dependence in C1-S1pr1, C2-ISG and C6-Cycling cells. These patterns indicate that perturbation responses learned from human cells were reshaped by the transcriptional programs already active in the target mouse cells. Because these TIL states differ in interferon activity, proliferative status and functional differentiation, the identity and magnitude of the response genes also varied across cellular contexts.

A small amount of target-species perturbation data further improved the prediction of state-specific responses. For *ARID1A*, Top20 Delta Pearson increased from 0.329 in the zero-shot setting to 0.839 and 0.862 after adaptation with 5% and 10% of the mouse perturbation cells, respectively. For *PDCD1*, the corresponding values increased from 0.421 to 0.760 and 0.764 (Fig. 5c and Supplementary Table S11). UMAP projections showed that, after few-shot adaptation, the predicted *ARID1A* and *PDCD1* response signatures were distributed more similarly to the observed mouse perturbation profiles across the TIL atlas (Fig. 5d and Supplementary Fig. S4). At the cell-state-aggregated level, 10% few-shot adaptation achieved a mean context Delta Pearson of 0.947 for both perturbations, with opposite-direction rates close to zero (Supplementary Table S12). These results show that a limited number of mouse perturbation samples can correct systematic cross-species differences while retaining the major response patterns learned from human data.

State-specific response genes further revealed distinct functional programs across TIL contexts. In C1-S1pr1 cells, *PDCD1*-associated response genes were enriched in IL-7 signaling, interferon-γ response, adaptive immune response and IL-6/JAK/STAT3 signaling (Fig. 5e). Genes including *PDCD1, STAT3, IL7R, CISH* and *ITGA4* were connected to these functional modules, linking the predicted response to cytokine signaling, T-cell homeostasis and immune regulation. In C2-ISG cells, *ARID1A* response genes were associated with cytokine signaling, TNF/NF-κB signaling, IL-2/STAT5 signaling and interferon-γ response, suggesting that perturbation of a chromatin-remodelling regulator may interact with inflammatory and cytokine-response programs in ISG-high cells (Supplementary Table S13). The sets of upregulated and downregulated genes induced by PDCD1 perturbation also differed among C1-S1pr1, C2-ISG, C3-41BB and C5-TermEx cells (Fig. 5f), showing that its transcriptional consequences were constrained by cellular state. Additional analyses similarly identified distinct immune- and cytokine-related programs for *ARID1A* and *PDCD1* across mouse TIL states (Supplementary Fig. S5).

In C5-TermEx cells, the BaiZe-predicted post-perturbation transcriptome for *PDCD1* was projected into morphology space using the pretrained MorphDiff G2I model [39] (Fig. 5g). The resulting DNA, ER, RNA, AGP and Mito channels provided candidate morphological readouts linked to the predicted perturbation response in this TIL state. Because no experimentally paired microscopy images were available, these outputs were interpreted as transcriptome-derived candidate morphology projections, with no image-level reconstruction validation.

### Temporal prediction recovers time-associated transcriptional changes and cell-state organization

Temporal state prediction provides a test of whether a conditional state-transition model can project early cellular states towards later developmental stages. Hematopoietic stem and progenitor cells undergo continuous transcriptional remodelling during differentiation, accompanied by dynamic changes in chromatin accessibility. To evaluate whether BaiZe could use early molecular states to predict subsequent differentiation stages, we constructed a cross-donor prediction task using the multimodal single-cell time-course dataset associated with GSE305370, which provides megakaryocyte and erythroid induction signatures [48]. The model learned state transitions from Day 2 to Day 3, Day 7 and Day 10 using the training donors, while donor 13176 was held out entirely for testing. For this donor, the Day 2 RNA state was used as the starting point, and later RNA states were predicted under three settings: RNA-only, randomly shuffled ATAC context and ATAC context matched to the target time point.

In the held-out donor, time-matched chromatin-accessibility information consistently improved the accuracy of later-state prediction (Fig. 6a). At Day 3, Day 7 and Day 10, Top20 Delta Pearson increased from 0.484, 0.731 and 0.836 in the RNA-only setting to 0.596, 0.785 and 0.897 with correct-time ATAC, respectively. The corresponding opposite-direction rates decreased from 0.353, 0.203 and 0.106 to 0.238, 0.117 and 0.035 (Supplementary Table S14). Shuffled ATAC produced results similar to RNA-only, indicating that the improvement depended on temporal correspondence between chromatin state and the target differentiation stage. Thus, the model benefited from chromatin context that captured stage-specific regulatory information.

**Figure 6.**
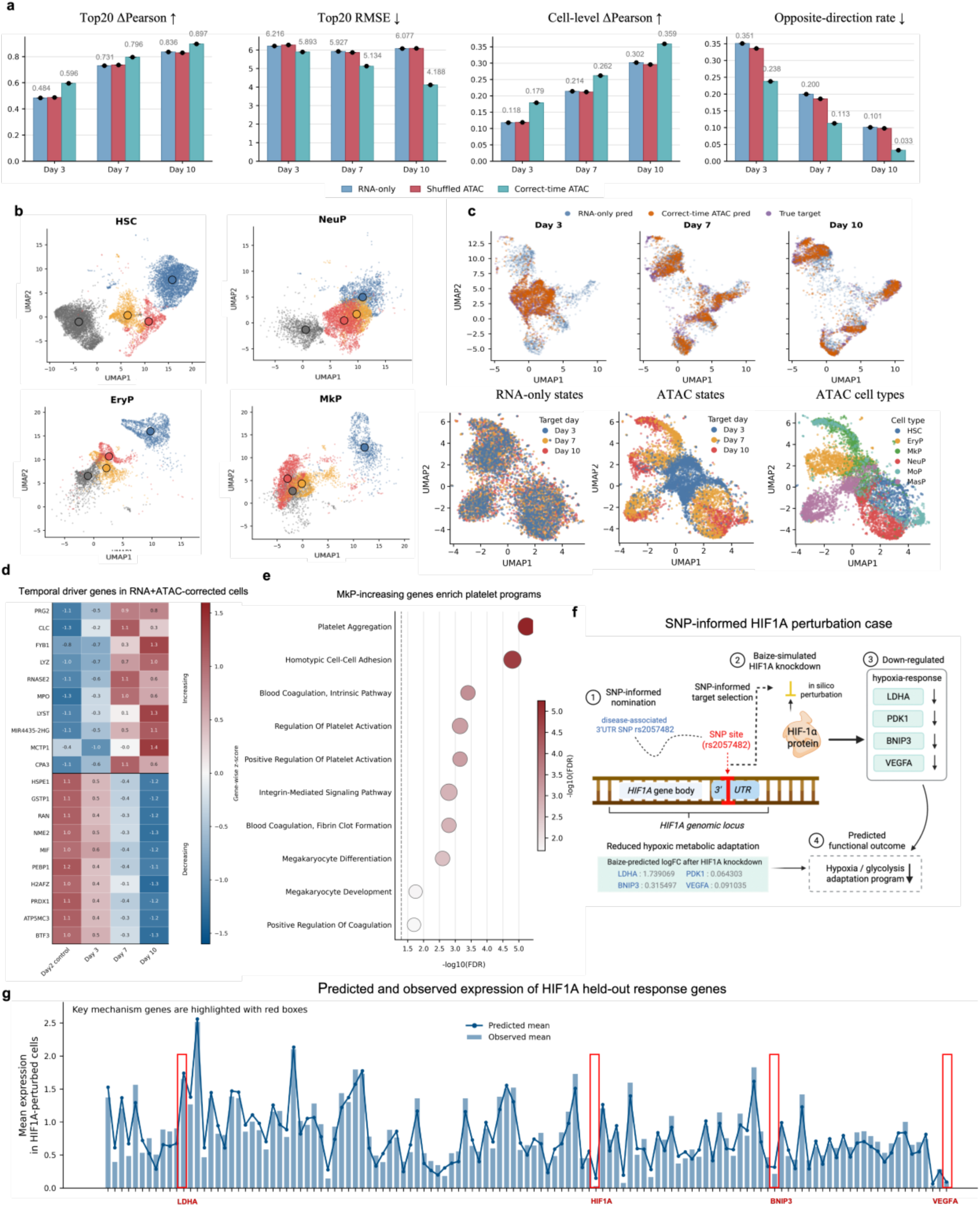
Temporal prediction and SNP-informed *HIF1A* perturbation modelling by BaiZe. (a) comparing RNA-only, shuffled ATAC and correct-time ATAC settings for Day3, Day7 and Day10 prediction in held-out donor 13176; (b) visualizing time-dependent ATAC accessibility changes across HSPC cell types from Day2 to Day10; (c) projecting predicted and observed RNA states onto shared UMAP spaces to assess recovery of temporal states and cell-type organization; (d) identifying temporal driver genes recovered in RNA+ATAC-predicted cells, including increasing and decreasing expression programs; (e) enriching MkP-increasing temporal genes for platelet activation, megakaryocyte differentiation and blood coagulation programs; (f) using rs2057482 as an SNP-informed clue to nominate *HIF1A* and simulate downstream responses after *HIF1A* inhibition; (g) comparing predicted and observed expression of held-out *HIF1A* response genes, with HIF1A-axis mechanism genes highlighted.

Chromatin-accessibility and transcriptomic embeddings further showed that time-matched ATAC information improved the recovery of differentiation-state structure. In the ATAC accessibility space, HSC, NeuP, EryP and MkP cells displayed varying degrees of displacement across Day 2, Day 3, Day 7 and Day 10, with the centers of the time-point distributions shifting along the differentiation process. This pattern indicates that chromatin states contain regulatory differences associated with temporal progression (Fig. 6b). In the RNA expression space, RNA-only predictions deviated from the observed target cells in several regions, whereas predictions generated with time-matched ATAC context showed greater overlap with the observed Day 3, Day 7 and Day 10 cell distributions (Fig. 6c). When colored by target day, the RNA-only predictions showed substantial mixing among temporal states. After incorporating time-matched ATAC information, Day 3, Day 7 and Day 10 cells occupied more distinct positions along the major branches of the embedding. These branches also corresponded to HSC, EryP, MkP, NeuP, MoP and MasP populations, indicating that the predictions recovered temporal differences while preserving the main lineage structure. Cell-type-specific RNA-state visualizations and joint projections of Day 2 cells with correct-time ATAC-predicted states showed consistent temporal shifts and lineage organization (Supplementary Fig. S6a,b).

The temporal genes recovered by BaiZe followed expression programs consistent with hematopoietic lineage progression (Fig. 6d). Genes including *PRG2, CLC, RNASE2, LYZ* and *MPO* increased over time and were associated with granulocytic, myeloid and immune-cell differentiation. In contrast, *HSPE1, GSTP1, RAN, NME2, PRDX1* and *ATP5MC3* gradually decreased, reflecting attenuation of metabolic, stress-response and basal cellular programs that were more active at earlier stages. Enrichment analysis of shared decreasing genes likewise indicated reduced proliferative, metabolic and RNA-processing activity during HSPC progression (Supplementary Fig. S6c). In MkP cells, genes that increased over time were enriched in platelet aggregation, platelet activation, megakaryocyte differentiation, integrin-mediated signaling and blood coagulation (Fig. 6e). These functions are consistent with megakaryocyte maturation, platelet formation and establishment of coagulation-related activity, showing that the predicted temporal changes retained biologically meaningful, lineage-specific developmental programs.

### SNP-informed perturbation modelling links HIF1A to downstream hypoxia-response programs

Disease-associated non-coding variants may influence cellular states by altering gene regulation and can therefore provide genetic clues for prioritizing candidate regulatory genes. To evaluate whether BaiZe could use such clues to infer downstream transcriptional responses, we selected rs2057482, a functional variant located in the 3′ untranslated region of *HIF1A* and near a miR-199a-binding site, as a genetic clue for nominating *HIF1A* as the perturbation target [49, 50]. *HIF1A* was treated as the candidate perturbation target, and CRISPR interference (CRISPRi) was used to approximate the cellular consequences of reduced *HIF1A* activity (Fig. 6f). This analysis did not model the allele-specific effect of rs2057482 or directly determine how the variant regulates *HIF1A*.

*HIF1A* is a central transcriptional regulator of cellular hypoxia responses and metabolic adaptation. BaiZe predicted that *HIF1A* inhibition would reduce the responses of hypoxia- and glycolysis-related genes, including *LDHA, PDK1, BNIP3* and *VEGFA* (Fig. 6f). LDHA supports the conversion of pyruvate to lactate, *PDK1* limits mitochondrial oxidative metabolism by inhibiting pyruvate dehydrogenase, *BNIP3* participates in hypoxia-associated mitochondrial homeostasis and cell-survival regulation, and *VEGFA* contributes to angiogenic signaling under low-oxygen conditions[51]. Their coordinated changes suggest that reduced *HIF1A* activity may weaken glycolytic adaptation, mitochondrial stress responses and broader cellular adaptation to hypoxia, thereby connecting the genetic clue to a functionally coherent downstream response program.

We further evaluated this prediction using the Replogle K562 CRISPRi dataset [1]. The model first learned general perturbation-response patterns from other gene perturbations, was then adapted using 33 *HIF1A*-perturbed cells, and was evaluated on 97 independent held-out cells; the complete dataset composition and split are summarized in Supplementary Table S1. Across selected *HIF1A*-related mechanism genes, the correlation between predicted and observed responses reached 0.893, and BaiZe recovered the observed directions of change for *HIF1A, LDHA, PDK1* and *BNIP3* (Fig. 6g). These results show that BaiZe can recover downstream response trends associated with the known function of an SNP-nominated candidate gene and convert a genetic clue into a testable perturbation hypothesis.

### An interactive agent enables natural-language interpretation of perturbation prediction results

BaiZe produces multiple layers of perturbation evidence, including expression changes, response-gene rankings, pathway enrichment, ATAC-based regulatory attribution, state-specific cross-species responses and morphology projections. These results are usually stored across expression matrices, gene tables, enrichment outputs and visualization files, which can make joint exploration difficult for users without extensive programming or bioinformatics experience. We therefore developed BaiZe-Agent, an interactive prototype that provides a accessible natural-language interface for exploring previously generated BaiZe results. Users can specify a perturbation, cellular context, drug dose, time point or species setting and receive a structured response linked to the corresponding evidence sources (Fig. 7a).

**Figure 7.**
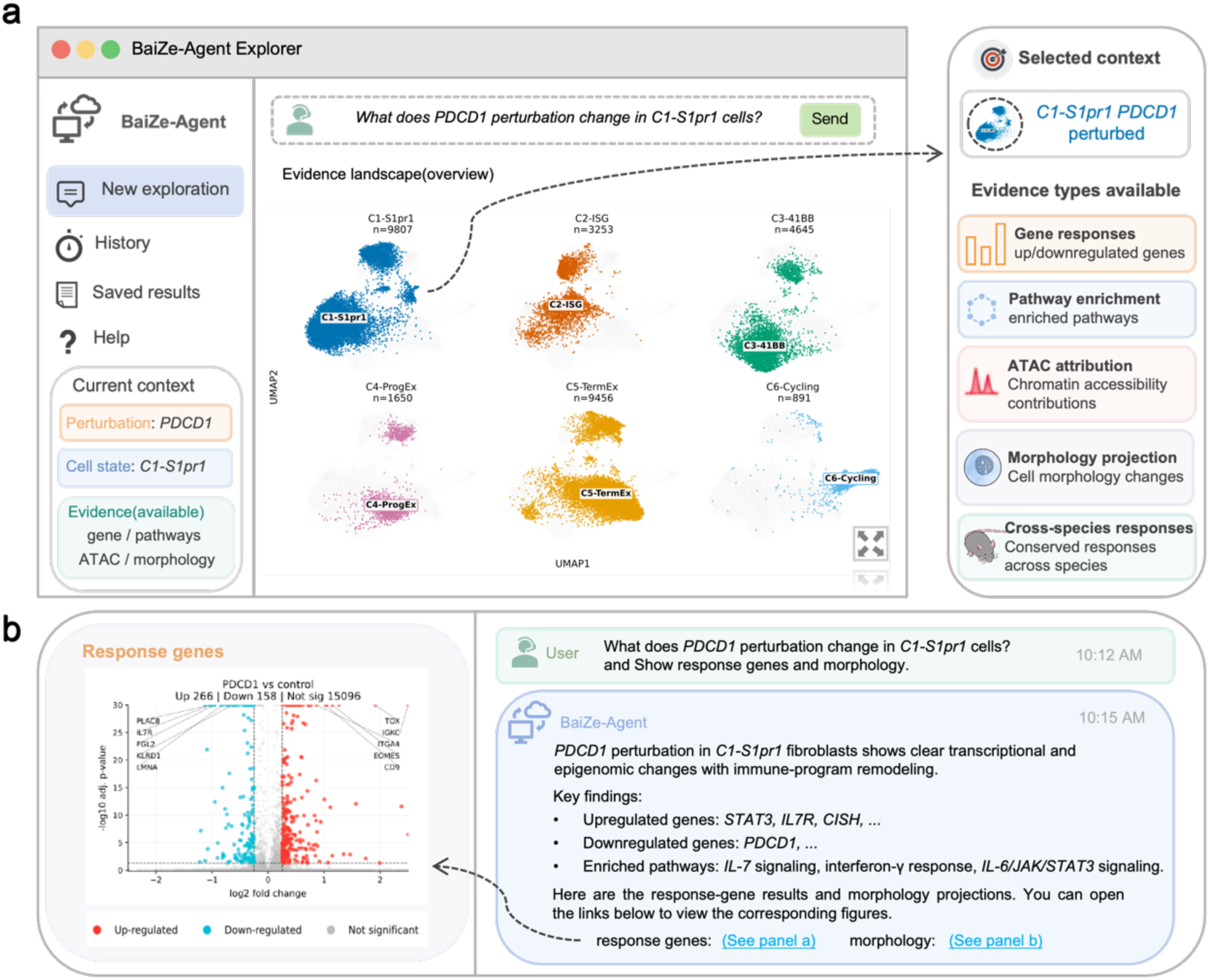
BaiZe-Agent enables natural-language exploration of perturbation predictions. (a) Exploration interface for selecting perturbation context and available evidence; (b) Example BaiZe-Agent response with links to supporting result views.

BaiZe-Agent retrieves information from saved BaiZe predictions and downstream analysis outputs. It identifies the biological context described in the query, selects the relevant evidence types and organizes the retrieved results into a concise response. Depending on the available outputs, the response may include major upregulated and downregulated genes, enriched pathways, ATAC-attributed regulatory regions, cross-species response patterns and morphology projections. Links to the corresponding tables, figures and analysis records are retained, allowing users to inspect the evidence underlying each response. Figure 7b illustrates this workflow using a representative perturbation query, in which the natural-language summary is connected to response-gene and morphology result views.

The interface is designed to support general result exploration across the perturbation settings evaluated in this study. In genetic and combinatorial perturbation tasks, it can retrieve response genes and functional programs. In drug-response and temporal analyses, it can organize results according to dose, cellular context or target time point. In ATAC-informed analyses, it can connect attributed chromatin regions with corrected genes and enriched pathways. In cross-species and morphology applications, it can retrieve state-specific responses and projected phenotype evidence. BaiZe-Agent operates on existing BaiZe outputs and does not perform additional model training, perturbation inference or statistical testing. Unsupported queries return an evidence-availability message, helping maintain traceability while reducing the technical barrier to exploring complex perturbation results.

## Discussion

BaiZe frames heterogeneous perturbation tasks as conditional cellular state transitions from a measured control state to a predicted post-perturbation transcriptome. Under this formulation, generalization draws on transferable response structure learned from perturbation data and cellular context represented by the control-state transcriptome. Their respective contributions are evident across held-out cell states, drug–cell-context shifts and human-to-mouse transfer. These results also highlight the importance of evaluating predictions in the expression-change space using complementary measures of correlation, magnitude error and response direction, because whole-transcriptome similarity can obscure errors among the most responsive genes.

The multi-view analyses indicate when additional molecular context may improve prediction and how the resulting evidence should be interpreted. Matched ATAC-seq context improved selected temporal-state predictions, whereas shuffled or masked accessibility features diminished this improvement, consistent with chromatin accessibility providing information complementary to baseline RNA [31]. Peak-level attribution identified accessibility features associated with model predictions, but these scores measure model dependence and do not establish causal peak-to-gene regulation. The cross-species results similarly support a source-to-target strategy in which human perturbation data provide an initial response estimate and limited mouse perturbation observations calibrate systematic differences [52,54]. Because the current analysis is restricted to one-to-one orthologs, species-specific genes and regulatory elements without direct correspondence are not represented.

Several BaiZe outputs are intended primarily to support hypothesis generation. Published associations involving rs2057482 were used to motivate the investigation of *HIF1A* [49, 50], and downstream transcriptional responses to CRISPRi-mediated *HIF1A* suppression were evaluated using the Replogle dataset [1]. MorphDiff maps predicted transcriptomes to candidate morphology projections [39], although the absence of paired microscopy prevents direct image-level validation. BaiZe-Agent facilitates access to stored response genes, pathways, ATAC-attribution results and morphology projections, but does not generate new perturbation predictions or statistical evidence. These distinctions define the evidential scope of the generated hypotheses.

The present study has several broader limitations. BaiZe was trained separately for the individual benchmarks, and the reported results therefore do not demonstrate a single universally pretrained model spanning all perturbation types and modalities. Performance remains sensitive to control matching, gene-space alignment, dataset-specific preprocessing and the availability of condition information, while point estimates do not fully characterize uncertainty in single-cell responses. Future work should investigate joint pretraining across perturbation datasets, calibrated predictive uncertainty, representations extending beyond one-to-one orthologs and prospective validation of predicted response genes, attributed regulatory regions and morphology projections. Within these limits, BaiZe provides a common modelling and evidence-organization framework for prioritizing context-dependent cellular-response hypotheses for experimental testing.

## Methods

### Datasets and preprocessing

We integrated publicly available single-cell perturbation and multi-omic datasets to evaluate BaiZe across genetic perturbation, combinatorial perturbation, drug response, ATAC-informed prediction, cross-species transfer, temporal state prediction and SNP-informed perturbation analysis. The Norman and Adamson Perturb-seq datasets [2, 4] were used for held-out double-gene combination prediction and prediction of the *ATF6+EIF2AK3+ERN1* triple-gene perturbation, respectively. Sci-Plex [5] was used for drug-response prediction conditioned on chemical structure and dose. Human embryo, MultiPerturb-seq and HSPC time-course multiome datasets [37, 43, 48] were used for unseen cell-state prediction, ATAC-informed modelling and cross-donor temporal prediction. Cross-species analysis used human CD8 T-cell perturbation data [54] as the source dataset and mouse tumour-infiltrating T-cell data [52] as the target system. Replogle K562 CRISPR interference data [1] were used for few-shot adaptation of *HIF1A* perturbation responses. Dataset modalities, biological systems, prediction tasks and split strategies are summarized in Supplementary Table S1.

All single-cell RNA datasets were organized as AnnData objects containing the expression matrix, gene identifiers, cell annotations and task-specific metadata. For raw count matrices, low-quality cells and lowly expressed genes were removed according to the original study or standard single-cell quality-control procedures. Counts were normalized by the total count of each cell and transformed using log1p:

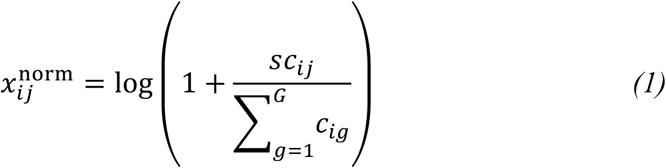

where *c_ij_* denotes the raw count of gene *j* in cell *i*, 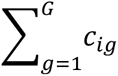 is the total count for cell *i*, *G* is the number of genes and *s* is the library-size scaling factor. When normalized expression matrices were provided by the original study, the processed values were retained without additional normalization. Gene identifiers, cell annotations and task-specific metadata were checked for consistency, and a fixed gene set and gene order were used within each task. Labels representing untreated or vehicle-treated cells, including control, ctrl, DMSO and vehicle, were standardized as control.

For tasks requiring an unperturbed starting state, control cells were matched according to cellular context. Controls with the same cell line, cell type or cell state, donor, time point and batch as the target cells were selected whenever available. When exact matching was unavailable, controls from the closest broader biological context were used, followed by the task-level control pool. Genetic perturbation labels were standardized to gene symbols, and multi-gene labels were parsed into target-gene sets, such as GENE1+GENE2 and GENE1+GENE2+GENE3. Drug metadata included drug name, SMILES representation, dose and cellular context.

ATAC-seq data were used as an optional chromatin-accessibility context. For paired RNA–ATAC datasets, the two modalities were matched at the single-cell level whenever possible; otherwise, matching was performed at the level of cell type, cell state, time point or another biological context. ATAC features were represented using low-dimensional embeddings supplied with the dataset or generated by the task-specific preprocessing pipeline. When single-cell ATAC embeddings were unavailable, context-specific embeddings were assigned from a preconstructed ATAC context bank. All ATAC features were aligned with the corresponding RNA samples by sample index and metadata before model input.

### Model architecture and residual diffusion objective

BaiZe formulates single-cell perturbation prediction as a conditional generation task from an unperturbed state to a post-perturbation state. Given an unperturbed RNA expression profile *x*_ctrl_ and condition information *c*, the model predicts the associated expression change and reconstructs the post-perturbation transcriptome. The observed expression change is defined as

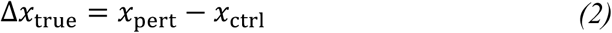

where *x*_pert_ denotes the observed post-perturbation RNA expression profile.

The model consists of a cellular-background encoder, a condition encoder, a deterministic response branch and a residual diffusion branch. The cellular-background encoder maps *x*_ctrl_ to a background representation *z*_bg_, and the condition encoder maps *c* to a condition representation *z*_c_. Given these representations, the deterministic branch estimates the principal expression change through a mean-delta head and a gene-prior head:

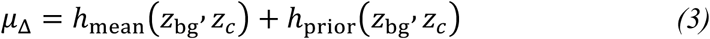

where ℎ_mean_ and ℎ_prior_ denote the mean-delta head and gene-prior head, respectively. The remaining component of the observed response is used as the residual target:

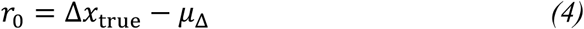

The residual diffusion branch applies a forward noising process to *r*_0_. For a randomly sampled timestep *t*,

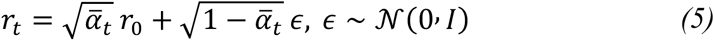

where 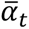 is the cumulative signal-retention coefficient at timestep *t*. The denoising network *f*_0_receives *r_t_*, *t*, *z*_bg_ and *z*_c_, and predicts the clean residual *r*_0_. The diffusion objective is

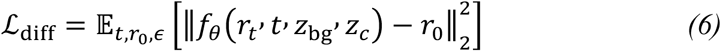

This formulation separates the perturbation response into a principal expression change estimated by the deterministic branch and a residual component modeled by diffusion.

During inference, the model first computes *μ*_Δ_ and then generates a residual estimate *r̂*_0_ through reverse diffusion. The predicted expression change is

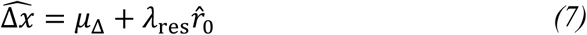

where *λ*_res_ is the residual sample scale. The predicted post-perturbation transcriptome is reconstructed as

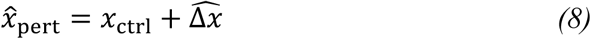

Model dimensions, diffusion timesteps, sampling steps and training configurations are summarized in Supplementary Table S2.

### Condition encoding for genetic, chemical, temporal and chromatin inputs

BaiZe maps task-specific intervention information into a shared condition representation. Given condition information *c*, the encoded condition is defined as

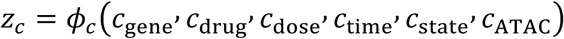

where *φ*_c_ denotes the condition encoder. Each task uses only the condition components available in that setting, and the resulting representation *z*_c_ is combined with the cellular-background representation *z*_bg_ during response prediction.

For genetic perturbation tasks, single-gene targets were standardized to gen e symbols, mapped to model indices and represented using learnable gene embeddings. Multi-gene perturbation labels, such as GENE1+GENE2 and GENE1+ GENE2+GENE3, were parsed into target-gene sets. The corresponding gene embeddings were encoded separately and combined into a fixed-dimensional condition representation. Control samples were assigned a dedicated control label.

For drug-response prediction, SMILES strings were converted into 2,048-bit Morgan fingerprints [53] and projected into the chemical-condition space. Treatment dose was encoded as a continuous variable and combined with the molecular representation. Drug names, known targets and mechanism-of-action annotations were retained for sample annotation and stratified analyses but were not used as predictive inputs. For temporal and state-transition tasks, the target time point or target cellular state was represented using a learnable embedding, while the input RNA profile defined the starting state. When ATAC-seq data were available, the low-dimensional chromatin-accessibility embedding was projected into the condition space and fused with the corresponding perturbation or temporal condition. The ATAC component was omitted in RNA-only models and included as an additional condition in RNA+ATAC models.

### Training strategy and inference procedure

BaiZe was trained separately for each benchmark using preprocessed RNA profiles, task-specific conditions and, where applicable, ATAC embeddings. Each dataset was divided into training, validation and test sets according to the task-specific split described in “Benchmark tasks and experimental settings.” Training samples paired an unperturbed RNA profile with the corresponding post-perturbation profile and condition information. Model parameters were optimized using the objective defined above, and validation performance was used to select the checkpoint for final evaluation. Post-perturbation profiles from the test set were excluded from model fitting, hyperparameter selection and checkpoint selection. Training configurations are summarized in Supplementary Table S2.

During inference, an unperturbed input profile was selected from the matched control pool and combined with the specified condition. BaiZe first estimated the deterministic expression change and then generated the residual component through reverse diffusion, yielding the predicted post-perturbation transcriptome. Predictions were retained at the single-cell level or aggregated according to the corresponding evaluation unit, including perturbation, cell type or state, drug–dose–cellular-context group, time point or species context. Aggregation procedures and evaluation metrics are described in “Evaluation metrics and statistical analysis.”

### Benchmark tasks and experimental settings

The benchmark suite evaluated generalization to held-out perturbations, unseen gene combinations and drugs, unseen cellular contexts and donors, and a target species. Test data were held out at the perturbation, drug, cellular-state, cell-line, donor or species level according to the corresponding task. Dataset-specific prediction tasks and split strategies are summarized in Supplementary Table S1.

Genetic perturbation benchmarks included held-out perturbation prediction in the Adamson dataset, held-out double-gene combination prediction in the Norman dataset, and prediction of the *ATF6+EIF2AK3+ERN1* triple-gene combination. In the human embryo task, the HYPO/VE state was excluded from training to evaluate prediction in an unseen cellular state. K562/RPE1 and MultiPerturb-seq benchmarks compared RNA-only and RNA+ATAC models, with shuffled, mismatched or region-masked ATAC controls used where applicable. The Sci-Plex benchmark included random cell, unseen-drug and held-out cell-line splits. For temporal prediction, Day 2 HSPCs were used as the starting state for prediction of Day 3, Day 7 and Day 10, with donor 13176 held out for cross-donor evaluation. The HIF1A task used a small number of *HIF1A* CRISPRi cells for adaptation and independent held-out cells for evaluation. Cross-species prediction was evaluated under zero-shot transfer and 5% or 10% target-species adaptation. For each benchmark, BaiZe was compared with task-appropriate perturbation-prediction models, pretrained single-cell models or statistical baselines using the same test cells, gene sets and evaluation metrics.

### Cross-species transfer and few-shot target-species adaptation

Cross-species analysis used human CD8 T-cell perturbation data as the source-species dataset and mouse tumour-infiltrating T-cell data as the target-species dataset. One-to-one human–mouse orthologs were obtained from Ensembl BioMart, yielding a shared space of 15,520 genes. Human and mouse expression matrices were reordered to an identical gene order, and all cross-species training, prediction and evaluation were performed in this shared gene space.

In the zero-shot transfer setting, BaiZe was trained using human CD8 T-cell perturbation data, while post-perturbation mouse RNA profiles were excluded from all model-fitting and selection procedures. During inference, the model received mouse control RNA from the target TIL state, the specified perturbation target and, where applicable, the corresponding mouse ATAC context. For few-shot target-species adaptation, the pretrained model was further adapted using either 5% or 10% of the mouse perturbation cells and evaluated on independent held-out mouse cells. Cells assigned to adaptation, validation and testing were kept separate, and mouse control cells were used only as unperturbed input backgrounds.

For each target TIL state, unperturbed inputs were preferentially selected from mouse control cells with the same state annotation. When state-matched controls were insufficient, a broader context-matched control pool was used. This matching strategy was used to limit variation arising from differences in target-cell state. Zero-shot, 5% few-shot and 10% few-shot results were reported separately. Because the analysis was restricted to one-to-one orthologs, the resulting predictions represent perturbation-response transfer within the shared gene space and do not include unmapped genes or other species-specific regulatory components.

### Evaluation metrics and statistical analysis

Model performance was evaluated primarily in the expression-change space. For the *i*-th test cell, the observed and predicted expression changes were defined as:

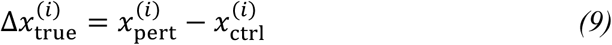

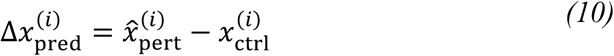

where 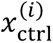 is the unperturbed input profile, 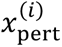 is the observed post-perturbation profile and 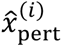 is the model prediction. Single-cell metrics were calculated from the expression-change vector of each test cell. For group-level evaluation, cells were aggregated by perturbation, cell type or cell state, drug–dose–cell context, time point or species context. The mean expression change for group *g* was defined as:

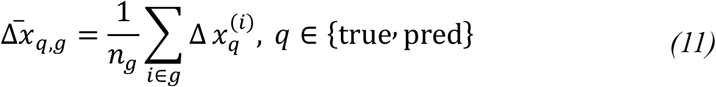

where *n*_g_ is the number of cells in group *g*. Group-level metrics assessed recovery of the average transcriptional response under a specified experimental condition.

Pearson correlation was used to measure agreement between predicted and observed expression changes:

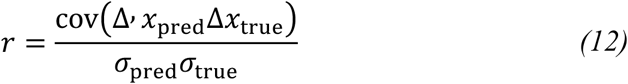

where *σ*_pred_ and *σ*_true_ are the standard deviations of the predicted and observed expression changes across genes. Mean squared error (MSE) was used to quantify errors in response magnitude:

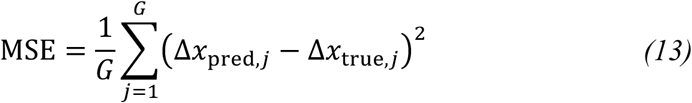

where *G* is the number of genes included in the evaluation. Both metrics were applied to either single-cell expression-change vectors or group-averaged response vectors.

To focus evaluation on genes with the largest observed responses, the Top *K*genes for each evaluation unit were selected according to the absolute observed expression change:

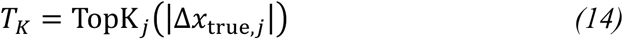

Top *K* Delta Pearson and Top *K* MSE used the same correlation and error definitions but were restricted to genes in *T _K_*. Unless otherwise stated, Top20 genes were used for genetic perturbation, temporal prediction and cross-species tasks, whereas Top50 genes were used in selected drug mechanism and dose analyses. Top *K*MSE was defined as:

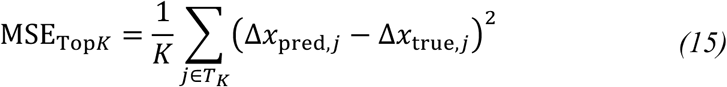

We further calculated the opposite-direction rate (ODR) to evaluate whether the regulation direction of major response genes was recovered correctly:

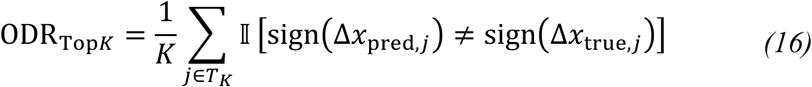

where II is the indicator function. A lower ODR indicates greater agreement in upregulation and downregulation, and direction accuracy was defined as 1 – ODR_TOP_*_K_*.

For experiments involving multiple random seeds or repeated data splits, we reported the mean across runs, with standard deviation or standard error reported where indicated in the corresponding figure or table. All methods were evaluated using the same test cells, gene sets and grouping definitions. Top *K* genes were selected exclusively from the observed test responses and were kept identical across models evaluated under the same test condition.

### ATAC-informed response correction and regulatory attribution

To identify response genes whose predictions were improved by chromatin-accessibility information, we compared gene-level absolute errors between the RNA-only and RNA+ATAC settings. For gene *j*, the ATAC-associated error reduction was defined as

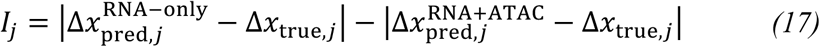

A positive *I_j_* indicates a lower prediction error after inclusion of ATAC context. Genes with *I_j_* > 0 and an RNA+ATAC prediction matching the observed response direction were defined as ATAC-corrected genes. Functional enrichment analysis was performed on these genes to identify biological processes and pathways associated with response correction.

To localize chromatin regions associated with corrected responses, ATAC features were ranked using their attribution scores and mapped back to the corresponding peaks. Peaks were annotated as promoter-proximal, gene-body or distal regions according to genomic position and were linked to the corresponding response genes and enriched functional programs. ATAC attribution scores quantify the contribution of individual accessibility features to model predictions; highly attributed peaks were interpreted as candidate regulatory evidence and require experimental validation before causal interpretation.

### Transcriptome-guided morphology projection

To assess whether BaiZe-predicted post-perturbation transcriptomes could support downstream phenotype projection, we used the pretrained MorphDiff gene-to-image (G2I) model to map transcriptomic states into five-channel Cell Painting morphology space [39]. MorphDiff uses a conditional latent diffusion process to generate DNA, endoplasmic reticulum (ER), RNA, actin/Golgi/plasma membrane (AGP) and mitochondrial (Mito) channels. The G2I checkpoint pretrained on the JUMP genetic perturbation dataset was used with all MorphDiff parameters frozen.

BaiZe-predicted post-perturbation transcriptomes were aligned by gene symbol to the fixed 12,328-gene MorphDiff input space and standardized using the MorphDiff reference expression distribution. Cross-species inputs were additionally clipped to the range from −4 to 4. Morphology projections were generated in G2I mode without control microscopy images. Gene-mapping coverage, sampling parameters and random seeds used for the displayed cases are summarized in Supplementary Table S15.

Because diffusion sampling is stochastic, multiple candidates were generated for each displayed case, and representative images were selected using predefined measures of sharpness, edge strength, contrast and saturation. Experimentally paired microscopy images were unavailable for the Adamson and mouse TIL settings. The resulting images were interpreted as transcriptome-derived candidate morphology projections, with no paired image data available for reconstruction-level validation.

### Natural-language querying with BaiZe-Agent

BaiZe-Agent was implemented as a web-based interface for querying, organizing and interpreting previously generated perturbation results. BaiZe-Agent v1.0 was implemented in Python 3.12 with Gradio 6.20.0 and a locally deployed Qwen3-8B model [55], which was used to convert retrieved evidence into concise natural-language explanations. During user interaction, the system did not retrain BaiZe or execute new perturbation-response inference. A rule-based query router identified the perturbation target, cellular context, drug–dose condition, target time point, species context and analysis intent, and matched these elements to task-specific records indexed in structured result files and an evidence registry. The retrieved records included saved prediction matrices, response-gene lists, pathway-enrichment results, ATAC-attribution outputs, cross-species analyses and morphology projections.

For matched queries, BaiZe-Agent organized the retrieved evidence into a structured natural-language response containing major upregulated and downregulated genes, associated pathways, candidate ATAC regions, state-specific or cross-species response patterns and available morphology-projection evidence. An evidence registry linked each response to the corresponding expression matrices, gene and pathway tables, attributed peaks, figures and source files, while deterministic routines handled numerical comparisons and other evidence-sensitive operations. The system did not perform new statistical analyses or extend biological conclusions beyond the retrieved evidence. When no matching record or sufficient supporting evidence was available, BaiZe-Agent returned an explicit evidence-availability message.

## Data availability

All datasets used in this study are publicly available through the original publications and public repositories. The human embryo 10x Multiome dataset used for unseen cell-state perturbation prediction was obtained from the Gene Expression Omnibus (GEO), accession number GSE218314 (https://www.ncbi.nlm.nih.gov/geo/query/acc.cgi?acc=GSE218314).The Norman combinatorial Perturb-se q dataset used for held-out multi-gene perturbation prediction was obtained fro m GEO, accession number GSE133344 (https://www.ncbi.nlm.nih.gov/geo/query/acc.cgi?acc=GSE133344).The Adamson UPR Perturb-seq dataset used for triple-gene perturbation extrapolation was obtained from GEO, accession number GSE 90546 (https://www.ncbi.nlm.nih.gov/geo/query/acc.cgi?acc=GSE90546).

The Sci-Plex drug perturbation dataset used for chemical structure- and dos e-conditioned response prediction was obtained from GEO, accession number G SE139944 (https://www.ncbi.nlm.nih.gov/geo/query/acc.cgi?acc=GSE139944). The MultiPerturb-seq RNA+ATAC dataset used for ATAC-guided perturbation predi ction, promoter peak masking and peak-level attribution analysis was obtained f rom GEO, accession number GSE277747 (https://www.ncbi.nlm.nih.gov/geo/query/acc.cgi?acc=GSE277747).

For cross-species perturbation prediction, the human CD8 T cell perturbation data were obtained from GEO, accession number GSE218988 (https://www.ncbi.nlm.nih.gov/geo/query/acc.cgi?acc=GSE218988). The mouse tumour-infiltrating T cell Perturb-seq and chromatin-accessibility data were obtained from GEO accession number GSE203593, including the ATAC-seq and Perturb-seq subseries. The HSPC time-course multiome dataset used for temporal RNA-state prediction was obtained from GEO, accession number GSE305370 (https://www.ncbi.nlm.nih.gov/geo/query/acc.cgi?acc=GSE305370).

The Replogle K562 CRISPRi Perturb-seq data used for the SNP-informed HIF1A perturbation case and related perturbation-response analyses were obtained from the processed Perturb-seq release on Figshare (https://plus.figshare.com/articles/dataset/_Mapping_information-rich_genotype-phenotype_landscapes_with_genome-scale_Perturb-seq_Replogle_et_al_2022_processed_Perturb-seq_datasets/20029387) and from the Genome-Wide Perturb-Seq portal (https://gwps.wi.mit.edu/). Human–mouse one-to-one ortholog information and gene annotations used for cross-species alignment and peak-to-gene annotation were obtained from Ensembl BioMart (https://www.ensembl.org/biomart/martview). Genome coordinates and reference genome annotations used for genomic-region annotation were obtained from the UCSC Genome Browser (https://genome.ucsc.edu/).

Gene-set and pathway resources used for enrichment analyses were obtained from the Molecular Signatures Database (MSigDB; https://www.gsea-msigdb.org/gsea/msigdb), Reactome (https://reactome.org/) and Enrichr (https://maayanlab.cloud/Enrichr/). The processed datasets, task-specific split files, prediction outpu ts, response-gene tables, enrichment results and ATAC attribution outputs gener ated in this study are available at GitHub (https://github.com/simonzqw/BaiZe).

## Code availability

The source code of BaiZe, including model training, inference, evaluation and downstream analysis scripts, is available at GitHub (https://github.com/simonzqw/BaiZe).

## Conflict of interest

The authors declare that they have no competing interests.

## Funding

This work was supported by the National Natural Science Foundation of China (No. 62476087, No. 62201341, and No. 82571771), the Shanghai Municipal Education Commission Initiative on Artificial Intelligence-Driven Reform of Scientific Research Paradigms and Empowerment of Discipline Leapfrogging, the National Key Research and Development Program of China (No. 2022YFB3203500), and the Natural Science Foundation of Shanghai (No. 25ZR1401167).

## Notes

https://www.ncbi.nlm.nih.gov/geo/query/acc.cgi?acc=GSE218314

https://www.ncbi.nlm.nih.gov/geo/query/acc.cgi?acc=GSE133344

https://www.ncbi.nlm.nih.gov/geo/query/acc.cgi?acc=GSE90546

https://www.ncbi.nlm.nih.gov/geo/query/acc.cgi?acc=GSE139944

https://www.ncbi.nlm.nih.gov/geo/query/acc.cgi?acc=GSE277747

https://www.ncbi.nlm.nih.gov/geo/query/acc.cgi?acc=GSE218988

https://www.ncbi.nlm.nih.gov/geo/query/acc.cgi?acc=GSE305370

https://github.com/simonzqw/BaiZe

### Competing Interest Statement

The authors have declared no competing interest.

