## Supplementary Information for "BaiZe: A Multi-View Dynamic Framework for Simulating and Interpreting Cellular Responses Across Perturbation Contexts"

#### Tables

Table S1. Dataset summary and task-specific preprocessing used for Baize evaluation.

| Dataset / module | Modality | Species / system | Main task | Split / key setting |
|---|---|---|---|---|
| Human embryo 10x Multiome | RNA + ATAC | Human embryo cell states: AME, EXMC, HYPO/VE | Unseen cell-state perturbation prediction | HYPO/VE held out as unseen cell state; Top20 genes selected by \|true delta\| |
| K562/RPE1 cross-cell-line perturbation | RNA ± ATAC | K562 and RPE1 | Cross-cell-line perturbation transfer | K562->RPE1 and RPE1->K562 settings; RNA-only vs RNA+ATAC comparison |
| Norman combinatorial perturbation | scRNA-seq | Genetic perturbation benchmark | Held-out double-gene combination prediction | GEARS-style seen0, seen1 and seen2 split; Additive vs Latent/Add or residual-corrected strategies |
| Adamson UPR Perturb-seq | scRNA-seq | K562 UPR perturbation system | Triple-gene perturbation extrapolation | Control, single-gene and double-gene perturbations used for training; ATF6+EIF2AK3+ERN1 held out for testing |
| SciPlex drug perturbation | scRNA-seq + chemical features | A549, K562 and MCF7 | Drug/dose response prediction | IID random, OOD unseen-drug, held-out K562/A549/MCF7 settings; Morgan fingerprint + dose used as input |
| MultiPerturb-seq RNA+ATAC | Paired RNA + ATAC | Human perturbation multiome | ATAC-guided response prediction and attribution | Human ENSG genes; 25,217 genes and 256-dimensional ATAC LSI representation; promoter masking and peak attribution |
| Human CD8 T -> | RNA + ATAC/context | Human CD8 T source | Cross-species | 15,520 one-to-one human- |

| Dataset / module | Modality | Species / system | Main task | Split / key setting |
|---|---|---|---|---|
| mouse TIL cross-species transfer | | and mouse TIL target | ARID1A/PDCD1 response prediction | mouse orthologous genes; zero-shot and 5%/10% mouse few-shot adaptation |
| GSE305370 HSPC time-course multiome | RNA + ATAC | Human CD34+ HSPC time course | Temporal RNA-state prediction | Day2 control to Day3/Day7/Day10; donor 13176 held out; RNA-only, shuffled ATAC and correct-time ATAC |
| Replogle K562 CRISPRi HIF1A case | scRNA-seq | K562 CRISPRi | SNP-informed HIF1A perturbation modeling | 39,052 cells, 8,248 genes and 120 background perturbations; HIF1A cells split into 33 train, 33 validation and 97 held-out test cells |

Table S2. Training and model architecture details of Baize.

| Model setting | Main task | Main condition inputs | Hidden dims | Perturb dim | Time dim | Dose dim | Attention heads | Timesteps | Sample steps | Batch size | GPU |
|---|---|---|---|---|---|---|---|---|---|---|---|
| Baize RNA-only | Genetic perturbation prediction | Control RNA, perturbation label | 512, 512, 512 | 200 | / | / | 4 | 1000 | 50 | 512 | 1 × vGPU-32GB |
| Baize RNA+ATAC | ATAC-guided perturbation prediction | Control RNA, perturbation label, ATAC context | 512, 512, 512 | 200 | / | / | 4 | 1000 | 50 | 512 | 1 × vGPU-32GB |
| Baize drug-response | Chemical perturba | Control RNA, Morgan fingerprint, dose | 512, 512, 512 | 200 | / | 32 | 4 | 1000 | 100 | 128 | 1 × RTX 4090D |

| | | | | | | | | | | | |
|---|---|---|---|---|---|---|---|---|---|---|---|
| | tion predicti on | | | | | | | | | | |
| Baize temporal | Time/st ate predicti on | Day2 control RNA, target time/state, optional ATAC context | 512, 512, 512 | 200 | 128 | / | 4 | 1000 | 100 | 256 | 1 × vGPU-32GB |

Table S3. Ablation study of ATAC context, scGPT gene prior and diffusion residual.

Component comparison

| **Method** | **Scale** | **Single Pearson ↑** | **Single MSE ↓** | **Single ΔPearson ↑** | **Single Top20 ΔPearson ↑** | **Perturb ΔPearson ↑** | **Perturb Top20 ΔPearson ↑** |
|---|---|---|---|---|---|---|---|
| RNA-only | 0.0000 | 0.7441 | 0.1823 | 0.0092 | 0.2648 | 0.0113 | 0.2183 |
| RNA-only + gene prior | 0.0000 | 0.7442 | 0.1847 | -0.0142 | 0.2280 | -0.0171 | 0.1576 |
| ATAC | 0.0000 | 0.7577 | 0.1639 | 0.1752 | 0.5098 | 0.2154 | 0.5663 |
| ATAC + gene prior | 0.0000 | **0.7616** | 0.1610 | 0.2099 | 0.5920 | 0.2589 | 0.5994 |
| ATAC + gene prior + Diffusion Residual | 0.1500 | 0.7547 | **0.1718** | **0.2332** | **0.7205** | **0.2882** | **0.7770** |

Residual sample scale = 0.15

| **Method** | **Scale** | **Single Pearson ↑** | **Single MSE ↓** | **Single ΔPearson ↑** | **Single Top20 ΔPearson ↑** | **Perturb ΔPearson ↑** | **Perturb Top20 ΔPearson ↑** |
|---|---|---|---|---|---|---|---|
| RNA-only | 0.15 | 0.7263 | 0.2052 | -0.0188 | 0.3047 | -0.0224 | 0.2654 |
| RNA-only + gene prior | 0.15 | 0.7264 | 0.2094 | -0.0380 | 0.2407 | -0.0455 | 0.1793 |
| ATAC | 0.15 | 0.7504 | 0.1762 | 0.2012 | 0.6922 | 0.2483 | 0.7729 |
| **ATAC + gene prior** | **0.15** | **0.7547** | **0.1719** | **0.2330** | **0.7204** | **0.2880** | **0.7771** |

Table S4. Combinatorial perturbation prediction under GEARS-style held-out splits.

| Group | Method | n | Top20 Delta Pearson | Top20 MSE |
|---|---|---|---|---|
| seen0 | Additive | 6 | 0.0000 | 0.7096 |
| seen0 | Residual-corrected | 6 | 0.3843 | 0.6898 |
| seen1 | Additive | 25 | 0.7699 | 0.3547 |
| seen1 | Residual-corrected | 25 | 0.7841 | 0.3357 |
| seen2 | Additive | 14 | 0.9491 | 0.0674 |
| seen2 | Residual-corrected | 14 | 0.9550 | 0.0526 |

Table S5. Representative UPR response genes in the held-out ATF6+EIF2AK3+ERN1 triple-gene perturbation case.

| Gene | Control mean | Observed mean | Predicted mean | Observed delta | Predicted delta |
|---|---|---|---|---|---|
| SDF2L1 | 4.4752 | 2.6560 | 3.4305 | -1.8192 | -1.0447 |
| HSPA5 | 2.4796 | 0.8750 | 1.5343 | -1.6046 | -0.9453 |
| HERPUD1 | 1.9355 | 0.3878 | 0.9380 | -1.5477 | -0.9975 |
| DDIT3 | 1.9103 | 0.4035 | 0.9669 | -1.5069 | -0.9434 |
| HSPA8 | 1.0633 | 2.2205 | 1.8016 | 1.1572 | 0.7383 |
| MANF | 1.6803 | 0.7547 | 1.1192 | -0.9256 | -0.5611 |
| HSP90B1 | 2.1627 | 1.2389 | 1.5780 | -0.9238 | -0.5848 |
| DDIT4 | 1.2704 | 0.3290 | 0.6476 | -0.9414 | -0.6228 |
| TRIB3 | 1.4766 | 0.4390 | 0.8354 | -1.0376 | -0.6411 |
| DNAJB11 | 1.2630 | 0.6426 | 0.8093 | -0.6203 | -0.4537 |
| SELK | 1.5667 | 0.9827 | 1.1376 | -0.5839 | -0.4291 |
| XBP1 | 1.1493 | 0.6502 | 0.8374 | -0.4991 | -0.3119 |

Table S6. Drug perturbation benchmark summary across random, unseen-drug and held-out cell-context settings.

| Setting | Held-out cell line | n cells | n groups | Cell Top20 Delta Pearson | Group Top20 Delta Pearson | Group MSE | Group Opp. |
|---|---|---|---|---|---|---|---|
| IID random split | NA | 102496 | 2244 | 0.7396 | 0.6927 | 0.0167 | 0.3002 |
| OOD unseen-drug split | NA | 39914 | 2218 | 0.7904 | 0.7334 | 0.0218 | 0.2680 |
| CRISP-like K562 held-out | K562 | 2573 | 36 | 0.7675 | 0.6852 | 0.0220 | 0.4505 |
| CRISP-like A549 held-out | A549 | 2954 | 36 | 0.4141 | 0.2289 | 0.0078 | 0.3738 |
| CRISP-like MCF7 held-out | MCF7 | 6323 | 36 | -0.0396 | -0.1080 | 0.2157 | 0.5332 |

Table S7. ATAC ablation and promoter peak masking performance.

| Model / ablation | Cell Delta Pearson | Cell Top20 Delta Pearson | Perturb Delta Pearson | Perturb Top20 Delta Pearson | Perturb MSE | Perturb Opp. |
|---|---|---|---|---|---|---|
| RNA-only | 0.0173 | 0.2621 | 0.0069 | -0.2833 | 0.2193 | 0.4287 |
| RNA+ATAC | 0.2279 | 0.3865 | 0.4638 | 0.7180 | 0.0600 | 0.1380 |
| RNA+ATAC with promoter peaks masked | 0.1965 | 0.3415 | 0.3684 | 0.4519 | 0.0774 | 0.1907 |

Table S8. Representative ATAC-corrected perturbation cases.

| Perturbation | RNA Top20 Delta Pearson | ATAC Top20 Delta Pearson | Gain Delta Pearson | RNA MSE | ATAC MSE |
|---|---|---|---|---|---|
| SMARCA4 | -0.7229 | 0.9148 | 1.6377 | 0.7712 | 0.0591 |
| BAZ1B | -0.7832 | 0.8900 | 1.6732 | 0.3078 | 0.0049 |

Table S9. Genomic region composition of top ATAC-attributed regions.

| Perturbation | Promoter fraction | Gene-body fraction | Distal fraction |
|---|---|---|---|
| BAZ1B | 0.7296 | 0.1586 | 0.1118 |
| BMI1 | 0.7779 | 0.1505 | 0.0716 |
| HDAC1 | 0.7407 | 0.1484 | 0.1109 |
| MBD3 | 0.7588 | 0.1497 | 0.0916 |
| PRDM16 | 0.7573 | 0.1416 | 0.1011 |
| SMARCA4 | 0.7504 | 0.1488 | 0.1008 |
| SMARCB1 | 0.7480 | 0.1505 | 0.1015 |
| UCHL5 | 0.7191 | 0.1695 | 0.1114 |

Table S10. Cross-species prediction performance across zero-shot and few-shot settings.

| Method | Setting | ARID1A Delta Pearson | PDCD1 Delta Pearson | Mean Delta Pearson | Mean MSE |
|---|---|---|---|---|---|
| Source-delta baseline | Zero-shot | 0.3682 | 0.2550 | 0.3116 | 0.1028 |
| scGPT-GenePrior beta=0.95 | Zero-shot | 0.3673 | 0.2546 | 0.3109 | 0.1031 |
| Baize W/ RNA | Zero-shot | 0.4327 | 0.3746 | 0.4036 | 0.1030 |
| Baize W/ ATAC | Zero-shot | 0.4490 | 0.4207 | 0.4348 | 0.1020 |
| Baize W/ ATAC | 5% mouse few-shot | 0.8381 | 0.7610 | 0.7995 | 0.0685 |
| Baize W/ ATAC | 10% mouse few-shot | 0.8591 | 0.8262 | 0.8427 | 0.0723 |

Table S11. Gene-level cross-species Top20 performance for ARID1A and PDCD1.

| **Perturbation** | **Method** | **Top20 Delta Pearson** | **Top20 MSE** |
|---|---|---|---|
| ARID1A | Source-delta | 0.2677 | 0.1142 |
| ARID1A | Zero-shot | 0.3292 | 0.1200 |
| ARID1A | 5% few-shot | 0.8389 | 0.0759 |
| ARID1A | 10% few-shot | 0.8615 | 0.0776 |
| PDCD1 | Source-delta | 0.2497 | 0.0930 |
| PDCD1 | Zero-shot | 0.4207 | 0.0880 |
| PDCD1 | 5% few-shot | 0.7602 | 0.0613 |
| PDCD1 | 10% few-shot | 0.7644 | 0.0677 |

Table S12. Context-level cross-species prediction performance across mouse TIL states.

| **Setting** | **Perturbation** | **Context cells** | **Mean context Delta Pearson** | **Mean context MSE** | **Mean opposite direction** |
|---|---|---|---|---|---|
| Zero-shot | ARID1A | 4668 | 0.9228 | 0.0489 | 0.0083 |
| Zero-shot | PDCD1 | 6465 | 0.9334 | 0.0331 | 0.0167 |
| 5% few-shot | ARID1A | 4435 | 0.9426 | 0.0392 | 0.0000 |
| 5% few-shot | PDCD1 | 6141 | 0.9382 | 0.0287 | 0.0083 |
| 10% few-shot | ARID1A | 4201 | 0.9465 | 0.0374 | 0.0000 |
| 10% few-shot | PDCD1 | 5818 | 0.9468 | 0.0285 | 0.0083 |

Table S13. Representative enrichment results for cross-species response genes and temporal programs.

| **Gene set** | **Library** | **Term** | **Adjusted P-value** | **Representative genes** |
|---|---|---|---|---|
| ARID1A C2-ISG context | Reactome 2022 | Cytokine Signaling in | 8.58e-07 | ITGB1; SOCS3; CCL5; CASP3; |

| Gene set | Library | Term | Adjusted P-value | Representative genes |
|---|---|---|---|---|
| | | Immune System | | STAT3; LTB; VIM; SQSTM1; CCR2 |
| ARID1A C2-ISG context | MSigDB Hallmark 2020 | TNF-alpha Signaling via NF-kB | 1.62e-08 | SOCS3; DUSP2; KLF6; CCL5; BHLHE40; MXD1; SQSTM1 |
| ARID1A C2-ISG context | MSigDB Hallmark 2020 | IL-2/STAT5 Signaling | 4.04e-07 | KLF6; CASP3; ODC1; BHLHE40; LTB; MXD1 |
| ARID1A C2-ISG context | MSigDB Hallmark 2020 | Interferon Gamma Response | 2.70e-04 | SOCS3; CCL5; CASP3; STAT3 |
| EryP increasing temporal genes | Hallmark | Heme Metabolism | 4.47e-04 | SLC25A37; MINPP1; DAAM1; TFRC; MBOAT2; FOXO3; ANK1 |
| MasP increasing temporal genes | Reactome | Immune System | 2.11e-03 | CD53; ATP8B4; FYB1; YES1; PRKCB; NFATC3; RAB27A; CSF2RB; GAB2; MPO |

Table S14. Temporal prediction performance by target day in held-out donor 13176.

| Method | Day | n cells | Cell Top20 Delta Pearson | Opposite direction |
|---|---|---|---|---|
| RNA-only | Day3 | 2628 | 0.4835 | 0.3532 |
| RNA-only | Day7 | 2724 | 0.7309 | 0.2027 |
| RNA-only | Day10 | 3000 | 0.8357 | 0.1059 |
| Shuffled ATAC | Day3 | 2628 | 0.4858 | 0.3383 |
| Shuffled ATAC | Day7 | 2724 | 0.7349 | 0.1856 |
| Shuffled ATAC | Day10 | 3000 | 0.8253 | 0.0981 |
| Correct-time ATAC | Day3 | 2628 | 0.5961 | 0.2377 |
| Correct-time ATAC | Day7 | 2724 | 0.7852 | 0.1172 |
| Correct-time ATAC | Day10 | 3000 | 0.8967 | 0.0352 |

Supplementary Table 15. MorphDiff configuration and inference settings for transcriptome-guided morphology projection.

| Category | Parameter | Setting |
|---|---|---|
| External model | Method | MorphDiff |
| External model | Inference mode | G2I |
| Input specification | Input dimension | 12,328 genes |
| Output specification | Output image size | 128 × 128 pixels |
| Output specification | Number of channels | 5 |
| Output specification | Channels | DNA, ER, RNA, AGP and Mito |
| Sampling | Sampling method | DDIM |
| Sampling | Sampling steps | 500 |
| Sampling | Guidance scale | 1.0 |
| Representative case | PDCD1 selected seed | 4210 |
| Representative case | ARID1A selected seed | 4304 |
| Gene mapping | Adamson mapped genes | 2,757 of 12,328 |
| Gene mapping | Cross-species mapped genes | 10,727 of 12,328 |
| Preprocessing | Cross-species clipping range | [−4, 4] |

### Figures

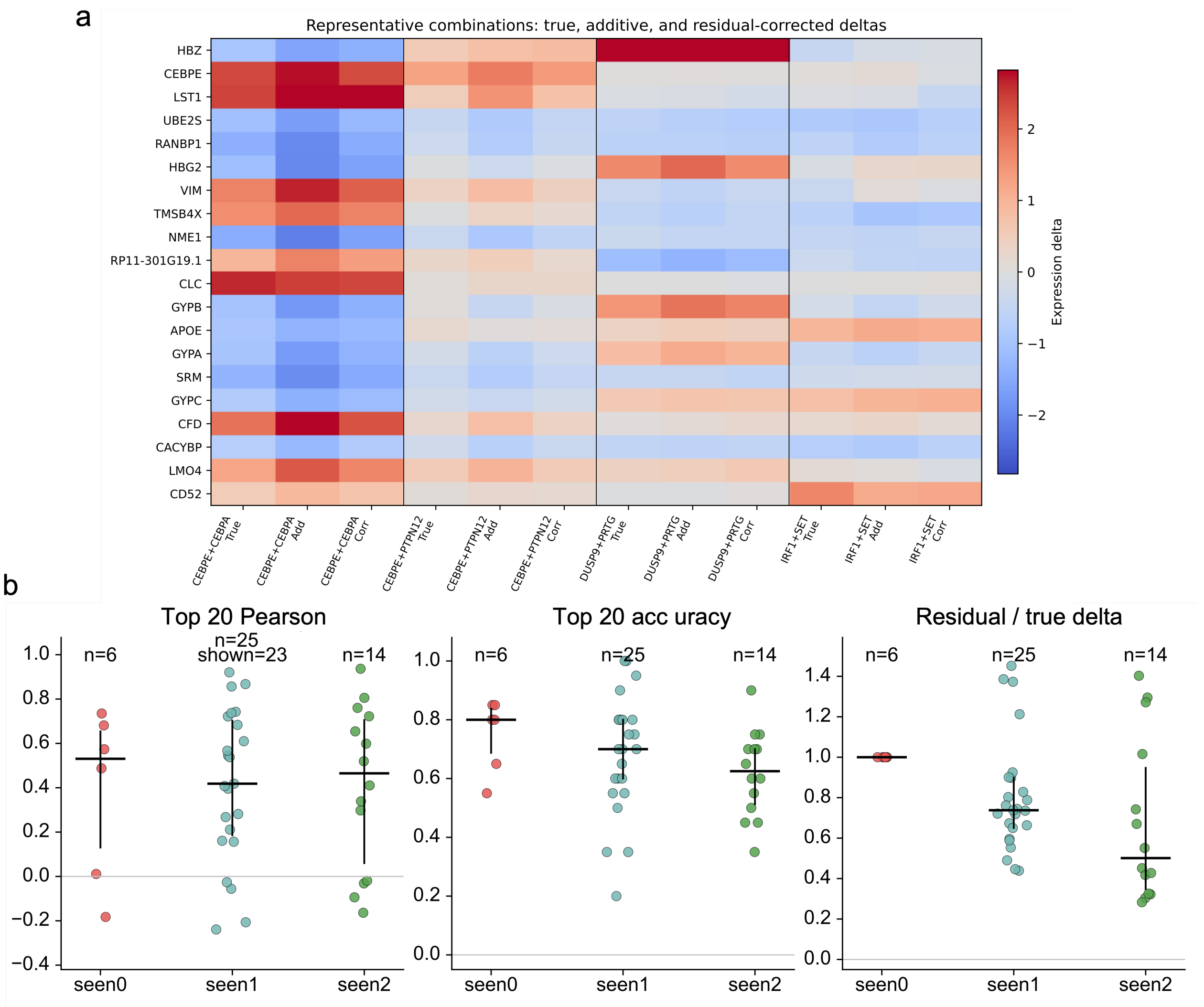


Supplementary Figure S1. Residual correction improves recovery of major response genes in held-out multi-gene perturbations.
(a) Representative heatmap of true, additive and residual-corrected deltas across selected double-gene perturbations.
(b) Distribution of Top20 Pearson, Top20 direction accuracy and relative residual magnitude across seen0, seen1 and seen2 combinations.

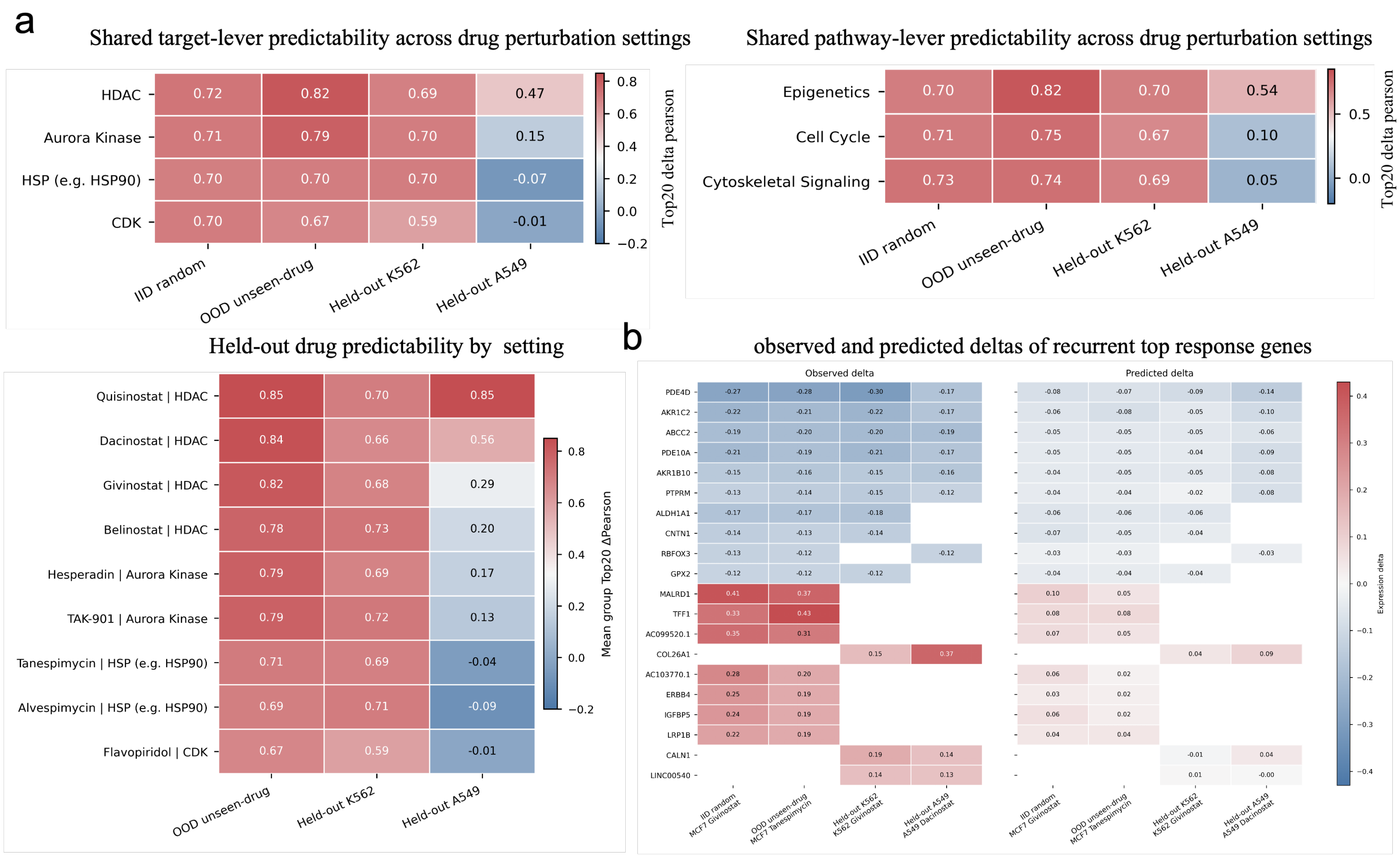


Supplementary Figure S2. Mechanism-level and recurrent-gene analysis of drug perturbation prediction.

(a) Heatmaps summarizing predictability across drug target classes, pathway classes and held-out drug settings. Values indicate Top20 or group-level Delta Pearson in IID random, OOD unseen-drug, held-out K562 and held-out A549 settings.

(b) Observed and predicted expression deltas of recurrent top response genes across representative drug perturbation settings. These analyses show that Baize recovers shared drug-response programs in several settings, while performance varies across drug mechanisms and held-out cell contexts.

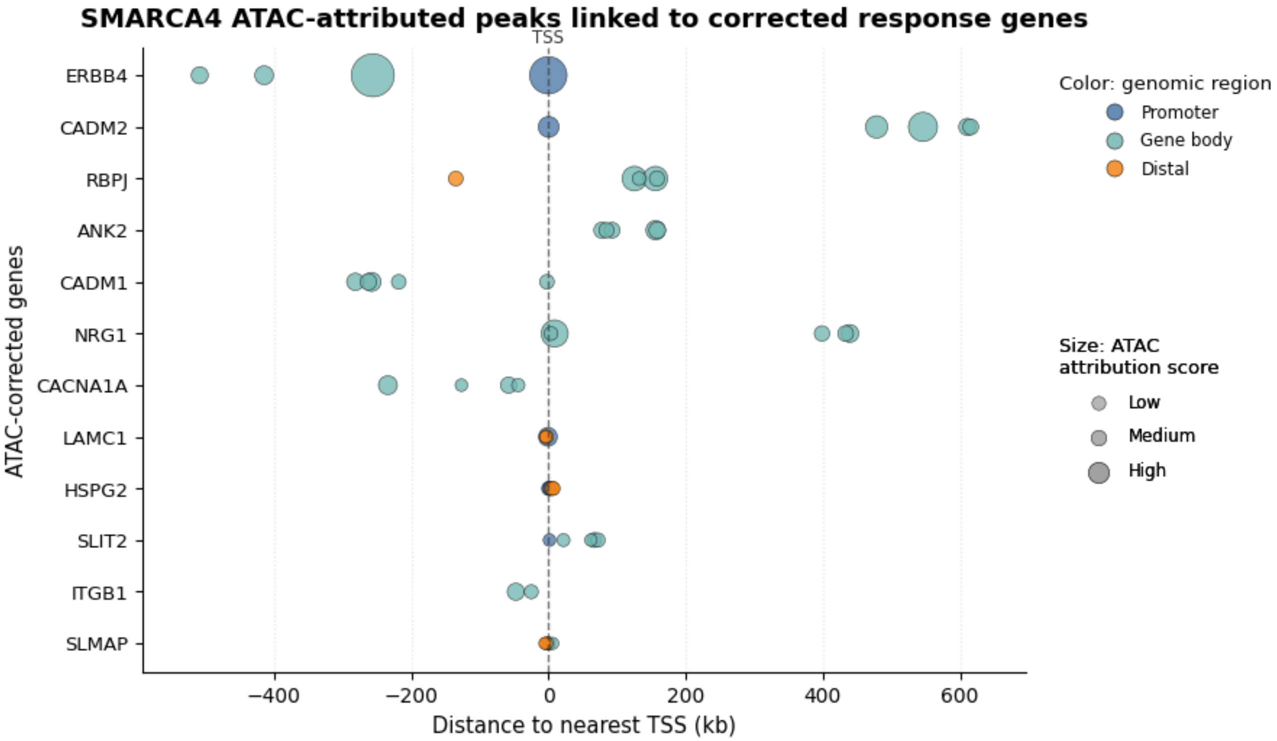


Supplementary Figure S3. Fine-grained SMARCA4 ATAC peak-to-gene attribution.

Each point represents an ATAC-attributed peak linked to an ATAC-corrected response gene under SMARCA4 perturbation. The x-axis shows the distance from the peak to the nearest transcription start site. Point color indicates genomic region class, and point size indicates ATAC attribution score. The dashed line marks the TSS.

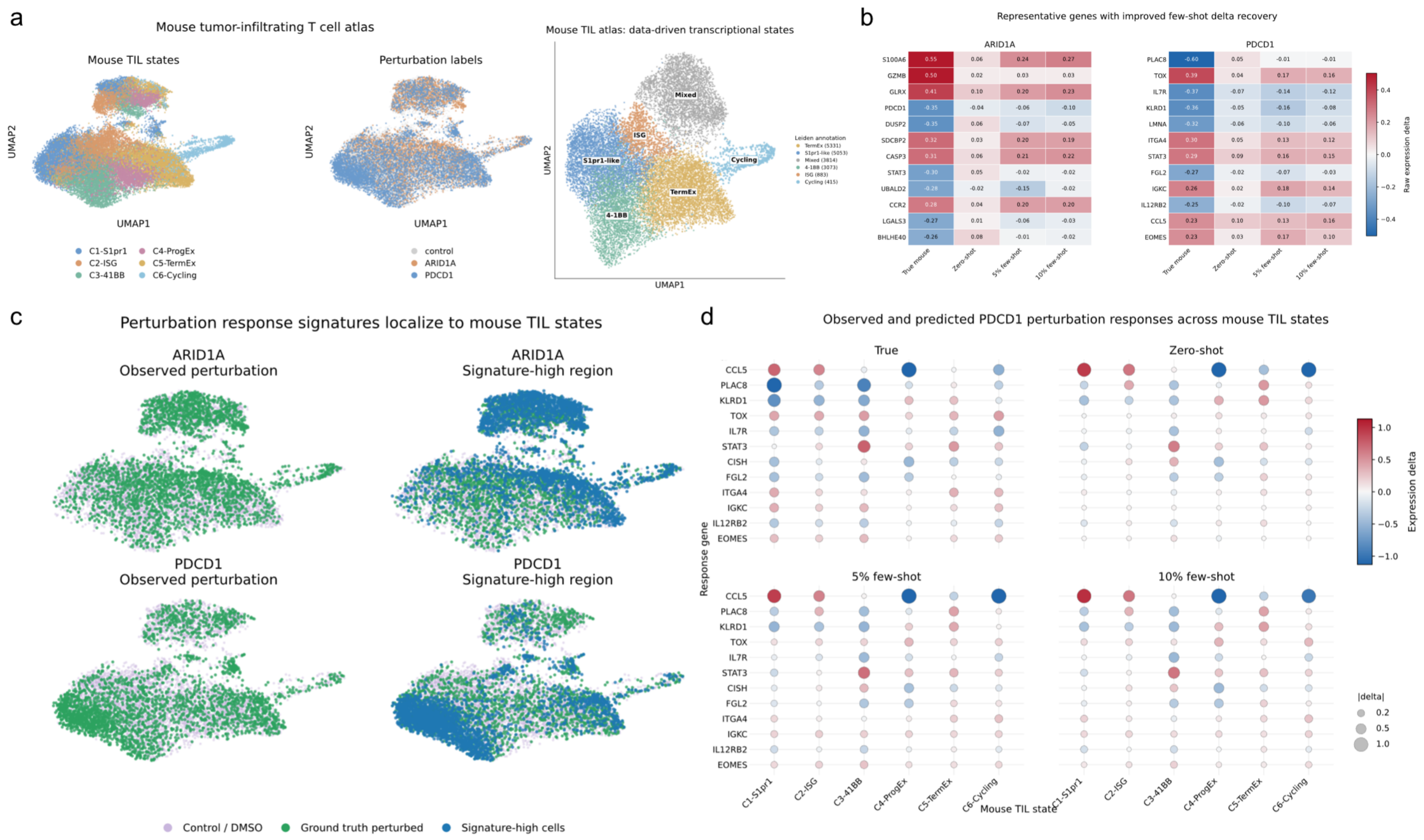


Supplementary Figure S4. Mouse TIL state structure and context-specific ARID1A/PDCD1 perturbation responses in cross-species prediction.
(a) UMAP visualization of the mouse tumor-infiltrating T cell atlas colored by annotated TIL states, perturbation labels and data-driven transcriptional states. The target mouse atlas contains six TIL contexts, including C1-S1pr1, C2-ISG, C3-41BB, C4-ProgEx, C5-TermEx and C6-Cycling.
(b) Heatmaps showing representative ARID1A and PDCD1 response genes with improved delta recovery after target-species few-shot adaptation. True mouse expression deltas are compared with zero-shot, 5% few-shot and 10% few-shot predictions.
(c) UMAP projections comparing observed ARID1A or PDCD1 perturbed cells with cells showing high perturbation-response signature scores. Perturbation signatures localize to specific regions of the mouse TIL state space, indicating state-dependent response patterns.
(d) Context-specific PDCD1 response profiles across six mouse TIL states. Dot color represents expression delta, and dot size represents absolute delta magnitude. Compared with zero-shot prediction, few-shot adaptation better recovers state-dependent PDCD1 response patterns.

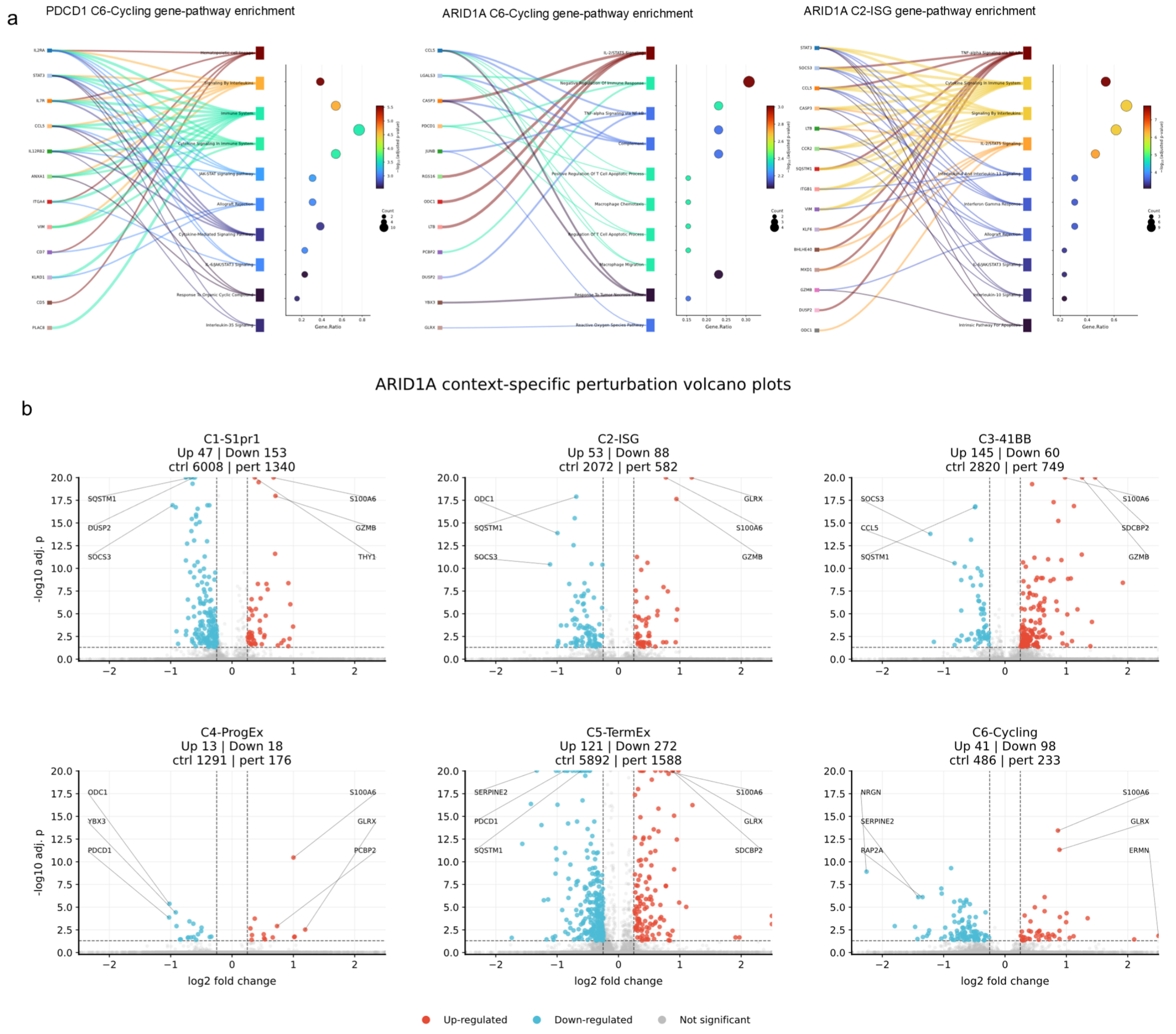


Supplementary Figure S5. Context-specific ARID1A/PDCD1 response programs in mouse TIL states.

(a) Gene-pathway enrichment plots for representative context-specific response genes from PDCD1 C6-Cycling, ARID1A C6-Cycling and ARID1A C2-ISG states. Links show gene-pathway associations, and dot plots indicate gene ratio, gene count and enrichment significance.

(b) Volcano plots showing ARID1A perturbation responses across six mouse TIL states. Red and blue points indicate significantly upregulated and downregulated genes, respectively. These results highlight heterogeneous, state-dependent ARID1A transcriptional responses.

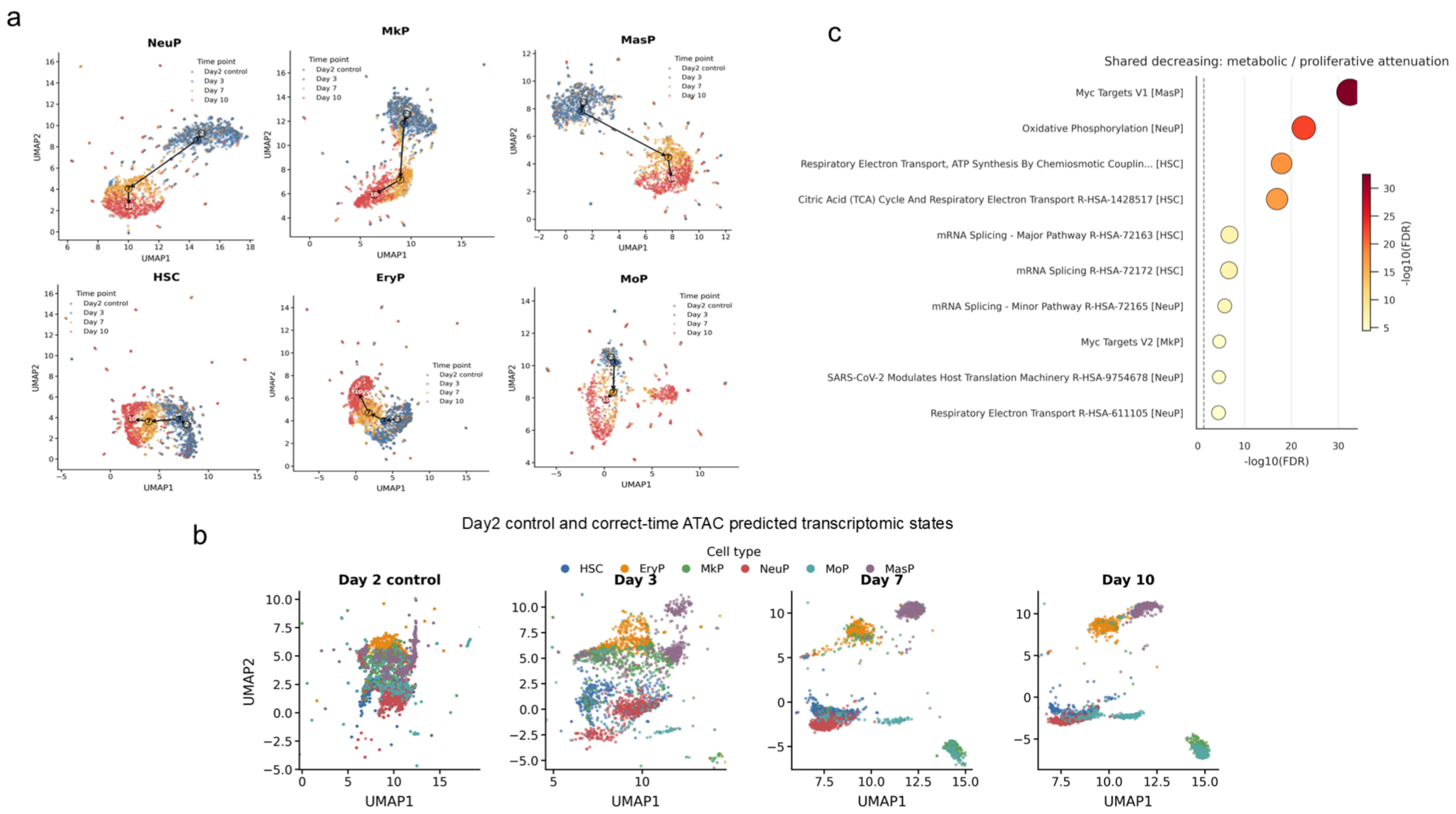


Supplementary Figure S6. Temporal transcriptomic-state organization and shared decreasing programs in HSPC time-course prediction

(a) UMAP visualization of RNA-derived states for six HSPC cell types across Day2 control, Day3, Day7 and Day10. Arrows indicate temporal shifts of transcriptomic states within each cell type.

(b) UMAP projection of Day2 control cells and correct-time ATAC-predicted transcriptomic states at Day3, Day7 and Day10, colored by cell type.

(c) Enrichment analysis of shared decreasing temporal genes, highlighting attenuation of proliferative, metabolic and RNA-processing programs during HSPC progression.
